# Leukocyte proliferation mediates disease pathogenesis in the *Ndufs4*(KO) mouse model of Leigh syndrome

**DOI:** 10.1101/2021.11.11.468271

**Authors:** Julia C Stokes, Rebecca L Bornstein, Katerina James, Kyung Yeon Park, Kira Spencer, Katie Vo, John C Snell, Brittany M Johnson, Philip G Morgan, Margaret M Sedensky, Nathan Baertsch, Simon C Johnson

**Author notes:** Authors contributed equally to this work (Julia C Stokes, Rebecca L Bornstein, Katerina James).

## Abstract

Symmetric, progressive, necrotizing lesions in the brainstem are a defining feature of Leigh syndrome (LS). A mechanistic understanding of the pathogenesis of these lesions has been elusive. Here, we report that leukocyte proliferation is causally involved in the pathogenesis of Leigh syndrome. Directly depleting leukocytes with a colony-stimulating factor 1 receptor (CSF1R) inhibitor dramatically attenuates disease, including complete prevention of CNS lesion formation and substantial extension of survival. Leukocyte depletion rescues a range of symptoms including hyperlactemia, seizures, respiratory function, and neurologic symptoms. These data provide a mechanistic explanation for the beneficial effects of mTOR inhibition. More importantly, these findings dramatically alter our understanding of the pathogenesis of LS, demonstrating that immune involvement directly drives disease. These findings have significant implication for the mechanisms of disease resulting from mitochondrial dysfunction, and may lead to novel therapeutic strategies.

**One-Sentence Summary:** Pharmacologic targeting of leukocytes prevents CNS lesions, neurological disease, and metabolic dysfunction in the *Ndufs4*(KO) mouse model of Leigh syndrome.

## Introduction

Considered as a group, genetic mitochondrial diseases (MDs) are the most common cause of heritable metabolic disease and one of the most common causes of pediatric neurological dysfunction (*1, 2*). MDs are genetically and clinically heterogeneous, with cases clustering into clinically defined syndromes (*2*). Subacute necrotizing encephalopathy, or Leigh syndrome (LS), is a multi-organ disease with metabolic, neurologic, and musculoskeletal symptoms, and is the most common form of pediatric MD (*3*). LS patients are often born healthy, showing symptom onset within the first few years of life. Symmetric progressive necrotizing lesions in the brainstem are a defining feature of LS, but a mechanistic understanding of these lesions has been elusive, and no effective interventions exist in the clinic. Inhibition of the mechanistic target of rapamycin (mTOR) attenuates disease in the *Ndufs4*(KO) mouse model of LS (*4, 5*), and mTOR inhibition appears to benefit some patients (*6, 7*), but the exact mechanisms underlying these benefits have been elusive.

Here, using small-molecule testing in the *Ndufs4*(KO) model, we report that the benefits of mTOR inhibitors can be attributed to inhibition of signaling mediated by the PI3K catalytic subunit gamma isoform, p110*γ*/PI3K*γ*, which is expressed mostly in leukocytes (*8*). We find that directly targeting leukocyte proliferation through inhibition of Colony Stimulating Factor 1 Receptor, CSF1R, dramatically attenuates disease in the *Ndufs4*(KO). CSF1R inhibition with the drug pexidartinib blocks CNS lesion formation, prevents neurologic symptoms, and extends survival. Strikingly, CSF1R inhibition also rescues symptoms not previously tied to CNS lesions or inflammation including hyperlactemia, seizures, hypoglycemia, and anesthetic responses. Critically, pexidartinib treated *Ndufs4*(KO) animals live as long as pexidartinib treated control mice, with drug toxicity, rather than MD, appearing to limit survival. Together, these findings provide evidence for the primary mechanism underlying the benefits of mTOR inhibition in LS and reveal that many symptoms of this disease have an immunologic origin amenable to direct pharmacologic targeting.

Note, while the accumulation of Iba1 (ionized calcium-binding adapter molecule 1, aka Allograft inflammatory factor 1/Aif1) (+) cells in LS lesions is typically referred to as microgliosis, here we attempt to avoid assigning cell type beyond what our staining specifically indicates, for reasons detailed in the discussion.

## Results

### *Isoform specific pharmacologic targeting of PI3Kγ significantly attenuates disease in the* Ndufs4(KO) *mouse model of LS*

To determine whether the benefits of mTOR inhibition in *Ndufs4*(KO) model result from disruption of PI3K mediated signaling, we treated animals from weaning (post-natal day 21, P21) with potent, orally available, isoform specific inhibitors of the catalytic subunits of PI3K: BYL719, GSK2636771, CAL-101, and IPI-549, inhibitors of p110α, p110β, p110δ, and p110γ, respectively.

Control treated *Ndufs4*(KO) animals display normal health early in life, but rapidly develop progressive neurological symptoms associated with CNS degeneration starting around P37, and death occurs by ∼P80 (***Fig. 1A***). Symptoms of neurologic decline include forelimb clasping, ataxia, and circling, presumed to result from the degenerative lesions in the brainstem and cerebellum. Additionally, animals develop cachexia without anorexia (***Fig. S1***); this is the most frequent proximal cause of death, as euthanasia due to loss of body mass is the approved study endpoint which is typically reached first (death by disease is generally not an allowed endpoint in IACUC approved animal work) (see ***Fig. 1***). The mechanistic underpinnings of this cachexia are unknown.

**Figure 1.**
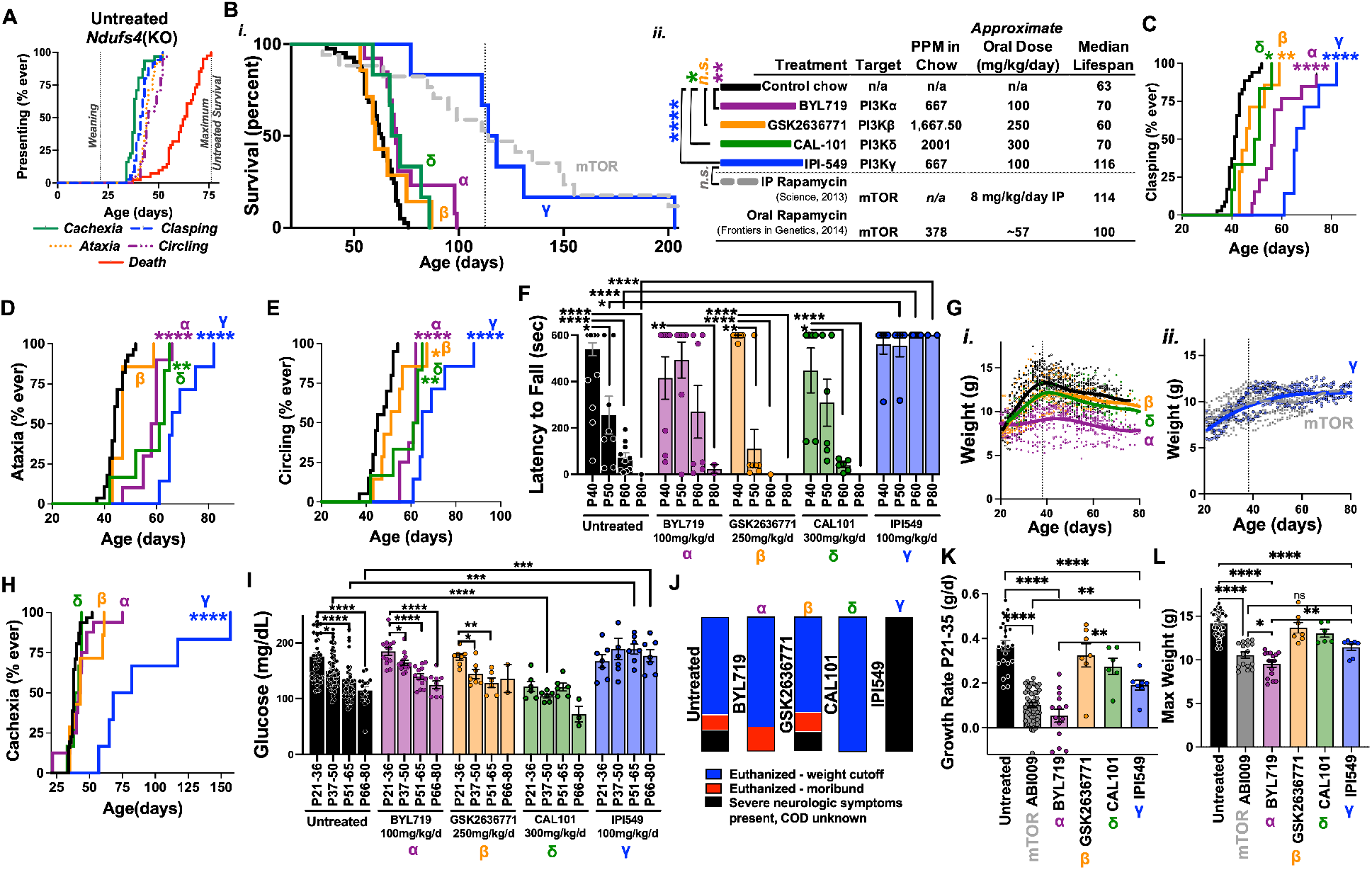
– Isoform specific inhibition of PI3K catalytic subunit p110*γ*, but not p110α, p110β, or p110δ, significantly attenuates disease in the *Ndufs4*(KO) mouse model of LS. (A) Disease course in untreated *Ndufs4*(KO) animals. Ages of death, onset of cachexia, and onset of observable behavioral symptoms related to neurologic dysfunction are shown. Disease symptoms on or onset shortly after postnatal day ∼37, with a rapid progression of symptoms until death by a median age of ∼P63 (in this study). Data here, and in subsequent graphs, are shown as ‘percent ever’ – when animals display a given symptom (see ***Methods***) they are scored and scoring for any given animal is not reversed even if the symptom is not observed on later dates. (B) Survival curves (i) and associated lifespan and dosage data (ii) for *Ndufs4*(KO) animals treated with isoform specific inhibitors of the PI3K catalytic subunits p110α, p110β, p110δ, or p110*γ* (BYL719 in purple, GSK2636771 in orange, CAL-101 in green, and IPI-549 in blue, respectively), or control chow (black). Published rapamycin treatment data is overlayed for reference (grey). Grey dashed line indicates median lifespan of rapamycin treated *Ndufs4*(KO) animals. (ii) Key for PI3K catalytic subunit inhibitor treatment lifespan curves with PPM in chow, approximate oral dosing, and median lifespans shown. Published rapamycin data provided for reference. *p<0.05, **p<0.005, and ***p<0.0005 by log-rank test. (C-E) Onset of clasping (C), ataxia (D), and circling (E) in *Ndufs4*(KO) mice treated with inhibitors of PI3K catalytic subunits p110α (purple), p110β (orange), p110δ (green), or p110*γ* (blue) (treatment groups also noted with Greek symbols above curves). Mice are scored when the symptom presents on at least two consecutive days. *p<0.05, **p<0.005, ***p<0.0005, and ****p<0.0001 by log-rank test vs untreated *Ndufs4*(KO) animals. (F) Performance of control and catalytic subunit specific inhibitor treated *Ndufs4*(KO) mice on a rotarod assay. *p<0.05, **p<0.005, ***p<0.0005, and ****p<0.0001 by unpaired, unequal variances (Welch’s), t-test. (G) Scatter plots of *Ndufs4*(KO) mouse weight as a function of age and treatment, with local regression (Lowess) curve overlayed to display population trends. (i) control, BYL719 (p110α inhibitor), GSK2636771 (p110β inhibitor), and CAL-101 (p110δ) treated *Ndufs4*(KO) mice. (ii) ABI-009 (mTOR inhibitor) and IPI-549 (p110*γ* inhibitor) treated *Ndufs4*(KO) mice. (H) Onset of cachexia in *Ndufs4*(KO) mice treated with inhibitors of PI3K catalytic subunits p110α (purple), p110β (orange), p110δ (green), or p110*γ* (blue) (treatment groups also noted with Greek symbols above curves). Cachexia onset, scored after completion of the survival studies, was scored as the day of life when an individual animal began showing weight-loss after ∼P37 (see ***Methods***). *p<0.05, **p<0.005, ***p<0.0005, and ****p<0.0001 by log-rank test vs untreated *Ndufs4*(KO) animals. (I) Blood glucose by age in control and PI3K catalytic subunit inhibitor treated *Ndufs4*(KO) animals. Each point represents the median value measured for one animal during the given time period (datapoints are biological replicates). *p<0.05, **p<0.005, ***p<0.0005, and ****p<0.0001 by unpaired, unequal variances (Welch’s) t-test. (J) Growth rate during the P21-P35 period of rapid post-natal growth. Datapoints represent individual animals. ****p<0.0001 by Welch’s ANOVA. *p<0.05, **p<0.005, ***p<0.0005, and ****p<0.0001 by unpaired, unequal variances (Welch’s) t-test. (K) Maximum animal weight over the course of lifespan. ****p<0.0001 by Welch’s ANOVA. *p<0.05, **p<0.005, ***p<0.0005, and ****p<0.0001 by unpaired, unequal variances (Welch’s) t-test. (L) Cause of death in survival studies by treatment group (see ***Methods***). For all panels, error bars represent SEM.

Treatment with the p110α, p110β, and p110δ inhibitors provided only modest (∼10% increase or less) benefits to survival. In contrast, treatment with the p110γ inhibitor IPI-549 increased survival similarly to mTOR inhibition – median survival in the IPI-549 treated *Ndufs4*(KO) mice was ∼110 days, versus ∼110 and ∼60 for rapamycin treated and untreated animals, respectively (***Fig. 1B***) (drug dosing was based on published studies (*9-12*); see ***Methods, Discussion***. With the exception of CAL-101, treatments led to similar tissue levels, ***Fig. S2***).

Each of the PI3K catalytic subunit inhibitors modestly delayed at least some behavioral symptoms of disease (clasping, circling, and ataxia): IPI-549 provided the greatest benefit, while BYL719 provided an intermediate benefit (***Fig. 1C-E***). In contrast, only IPI-549 improved *Ndufs4*(KO) performance on a rotarod assay, which assesses neurologic function and overall health (***Fig. 1F-G***); only IPI-549 impacted the onset of cachexia (***Fig. 1H***), and, as detailed, only IPI-549 increased survival. Additionally, IPI-549 alone prevented progressive hypoglycemia, which occurs during disease progression in the *Ndufs4*(KO) (***Fig. 1I***). IPI-549 alone qualitatively impacted cause of death: euthanasia resulting from body mass loss did not occur in IPI-549 treated animals, consistent with the rescue of cachexia in this group (***Fig. 1J***).

Rapamycin at doses sufficient to attenuate disease significantly reduces developmental weight gain and maximum body size (***Fig. 1K-L***) (*4, 5*). IPI-549 and mTOR inhibition similarly modify disease, while IPI-549 had a milder impact on growth (***Fig. 1K***). In contrast, BYL719 severely impaired growth and size, significantly more than rapamycin, while providing only modest benefits to disease (***Fig. 1B-L***). Together, these data indicate that mTOR inhibition does *not* benefit MD through actions on insulin/IGF-1 signaling (IIS) as p110α, not p110γ, mediates IIS (see ***Discussion***).

For thoroughness, we also tested a pan-PI3K inhibitor, BKM-120, but found that while well-tolerated in adult mice up to 60 mg/kg/day (*13*), BKM-120 was not tolerated at 50 or 100 mg/kg/day when started at weaning (***Fig. S3***). Given our IPI-549 results, we did not explore this strategy further.

### *mTOR inhibition reduces Iba1(+) leukocyte proliferation* in vitro

p110γ is primarily expressed in leukocytes (***Fig. S4***), which include the brain’s resident macrophages - microglia. Lesions in LS are characterized in part by the accumulation of Iba1(+) leukocytes (typically referred to as microgliosis, see discussion regarding use of ‘leukocyte’ versus ‘microglia’). This is widely thought to be a secondary reaction to CNS cell death caused by some combination of ‘energetic depletion’, reactive oxygen species (ROS) damage, lactic acidosis, and excitotoxicity (*3, 14*). Given that p110γ and mTOR inhibitors provide similar benefits, we reasoned that leukocyte (including microglia) proliferation may be a causal driver of CNS lesion formation and degeneration, rather than simply a secondary response.

To assess whether mTOR inhibition impacts leukocyte/microglia numbers, we tested the impact of ABI-009 (aka nab-rapamycin, see (*15*)), a water-soluble nano-particle formulation of rapamycin (see ***Methods***), on Iba1(+) cells in a mixed brain cell culture assay and compared the impact with that of pexidartinib/PLX3397, a colony stimulating factor 1 receptor (CSF1R) inhibitor which blocks leukocyte survival signaling (*16*) (***Fig. 2A***). ABI-009 and pexidartinib both reduced the fraction of Iba1(+) cells present in mixed neonatal brain-cell cultures in a dose-dependent manner, while the maximum effect of ABI-009 was only ∼50% of total compared to a near complete depletion of microglia with pexidartinib. Accordingly, mTOR inhibition does appear to preferentially limit leukocyte proliferation compared to other cell types, but the potency is reduced compared to direct targeting of leukocyte survival through CSF1R.

**Figure 2.**
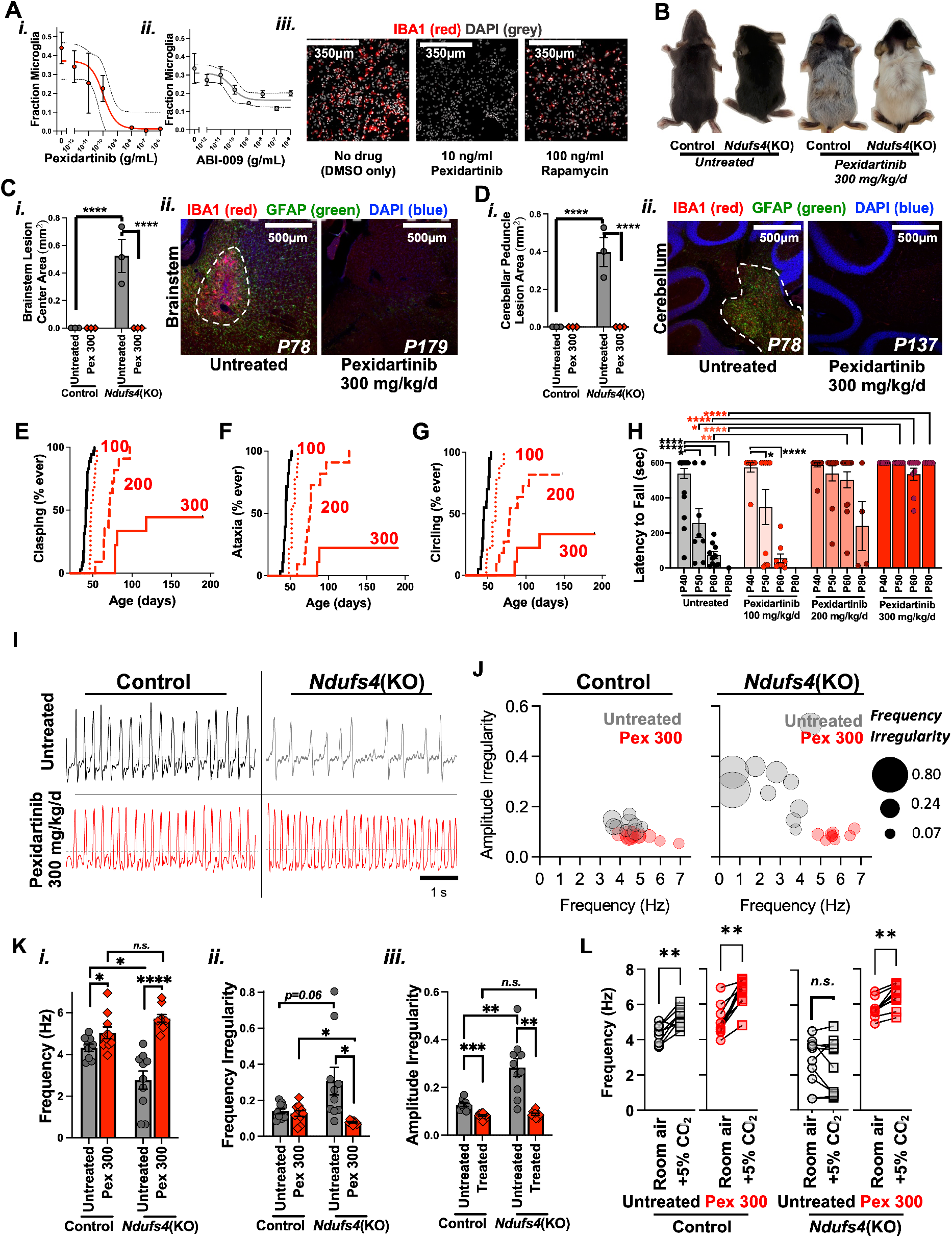
– Leukocyte depletion prevents CNS lesions and significantly attenuates disease in the *Ndufs4*(KO) model of LS. (A) Dose-dependent impact of the CSF1R inhibitor pexidartinib (i) and mTOR inhibitor rapamycin (ABI-009) (ii) on the fraction of Iba1(+) leukocytes (in this instance microglia due to source for cultures, see ***Methods***) in mixed primary brain cultures. Error bars represent SEM, dashed lines show the 95% confidence interval for an [Inhibitor] vs. response (three parameters) least squares fit. (iii) Representative images of mixed primary brain cultures stained with an anti-Iba1 antibody (green) and DAPI (blue, nuclei). (B) Representative pictures of control and *Ndufs4*(KO) animals treated with control diet or ∼300 mg/kg/day pexidartinib via chow. Pexidartinib treatment caused animal fur to whiten. (C-D) Quantification of brainstem (C) and cerebellar peduncle (D) lesion size (area of lesion in central slice in serial sectioning, see ***Methods***) in control and 300 mg/kg/d pexidartinib treated control and *Ndufs4*(KO) animals. Quantification (i) and representative images (ii) for both regions. Representative images are only provided for *Ndufs4*(KO) animals as control mice do not develop lesions (see ***Fig. 3*** for quantification of Iba1(+) leukocytes and GFAP(+) astrocyte cell numbers in control and *Ndufs4*(KO) mice). Anti-Iba1antibody staining in red, DAPI (nuclei) in greyscale (see ***Methods***). Lesion areas indicated by dashed white lines. Ages are noted in representative images (P78, etc). ****p<0.0001 by unpaired, unequal variances (Welch’s), t-test. (E-G) Onset of clasping (E), ataxia (F), and circling (G) in *Ndufs4*(KO) mice fed control diet (black lines) or administered pexidartinib at 100 (dotted red lines), 200 (dashed red lines), or 300 (solid red lines) mg/kg/day *Ndufs4*(KO) mice. *p<0.05, **p<0.005, ***p<0.0005, and ****p<0.0001 by log-rank test vs untreated *Ndufs4*(KO) animals. (H) Performance of control and pexidartinib treated animals on a rotarod assay. *p<0.05, **p<0.005, and ****p<0.0001 by unpaired, unequal variances (Welch’s), t-test. (I) Representative traces of breathing activity in P70 (+/- 2 days) control and *Ndufs4*(KO) mice fed control chow or administered 300 mg/kg/d pexidartinib. (J) Multivariable plotting of respiratory amplitude irregularity, frequency, and frequency irregularity in P70 (+/- 2 days) control and *Ndufs4*(KO) mice fed control chow or administered 300 mg/kg/d pexidartinib. (K) Single variable analysis of data in panel (J). Datapoints represent individual animals, error bars show SEM. *p<0.05, **p<0.005, and ****p<0.0001 by unpaired, unequal variances (Welch’s), t-test. (L) Respiratory responses to increased environmental CO2. Pairwise data shown for responses in individual mice **p<0.005 by Wilcoxon matched pairs signed rank test.

### Leukocyte depletion prevents CNS lesions and associated neurologic sequalae, including respiratory failure

Taken together, our data suggested that leukocyte proliferation may causally drive disease in LS. This possibility has not previously been explored and, if true, would dramatically alter our understanding of the pathogenesis of LS.

To test this model, we treated *Ndufs4*(KO) and control animals with 100, 200, or 300 mg/kg/day pexidartinib in normal mouse chow (dosing is approximated based on food consumption, see ***Methods***; brain and liver drug levels in ***Fig. S5***; a brief note - the higher doses led to a change in mouse coat color, consistent with reports of hair whitening in humans (***Fig. 2B***)).

Treatment with 300 mg/kg/day pexidartinib led to a complete (by IHC) prevention of brainstem (***Fig. 2C***) and cerebellar (***Fig. 2D***) lesions in *Ndufs4*(KO) mice, even at ages far beyond the maximum survival of untreated animals (survival data below). Importantly, both accumulation of Iba1(+) cells *and* astrocytosis were completely prevented by pexidartinib; the rescue of astrocytosis by a leukocyte inhibitor suggests that astrocyte involvement is secondary to leukocyte activity, a detail of LS pathobiology not previously resolved.

Consistent with these histological findings, pexidartinib dose-dependently delayed the onset and reduced overall incidence of behavioral signs of brainstem and cerebellar degeneration - forelimb clasping, ataxia, and circling (***Fig. 2E-G***). Pexidartinib treatment also rescued performance on the rotarod assay (***Fig. 2H***). Impaired respiratory center activity, which is a brainstem function, is a proximal cause of death in LS patients and has been reported to be a proximal cause of death in *Ndufs4*(KO) mice not euthanized due to weight criteria (*17*). To determine whether respiratory function is rescued by leukocyte depletion, we performed plethysmography in untreated and 300 mg/kg/day pexidartinib treated mice (see (*18*) for diagram, ***Methods*** for details). Severe defects in respiratory function were present in untreated *Ndufs4*(KO) mice at ∼P70, with defects in overall respiratory frequency, increased incidence of frequency and amplitude irregularities, and defective responses to increased CO_2_ (***Fig. 2I-L***). Remarkably, treatment with 300 mg/kg/day pexidartinib completely prevented each of these measures of respiratory dysfunction (***Fig. 2I-L***).

### Leukocyte depletion rescues neuroinflammation outside of CNS lesions

Overt lesions in the brainstem and cerebellum underlie many defining features of LS, but inflammation in other brain regions has not been carefully studied. Given the robust impact of pexidartinib in preventing lesions, we wondered whether neuroinflammation is present, and responsive to pexidartinib, in other *Ndufs4*(KO) brain regions. To probe this possibility, we analyzed Iba1(+) and GFAP (+) (astrocyte) cell numbers in cortex and in brainstem regions outside of overt lesions. Untreated *Ndufs4*(KO) mice show increases in Iba1(+) cells and astrocytes in non-lesion CNS tissue in the *Ndufs4*(KO); 300 mg/kg/day pexidartinib prevented both signs of neuroinflammation (***Fig. 3A-B***, see also ***Fig. S6***). Notably, pexidartinib treatment led to a near-complete depletion of Iba1(+) cells while astrocytes were rescued to control levels, again indicating that astrocytosis is secondary to leukocyte involvement. As with lesions, inflammation was prevented by pexidartinib even in mice far older than the maximum lifespan of untreated *Ndufs4*(KO)s (see ***Fig. 3*** legend for details).

**Figure 3.**
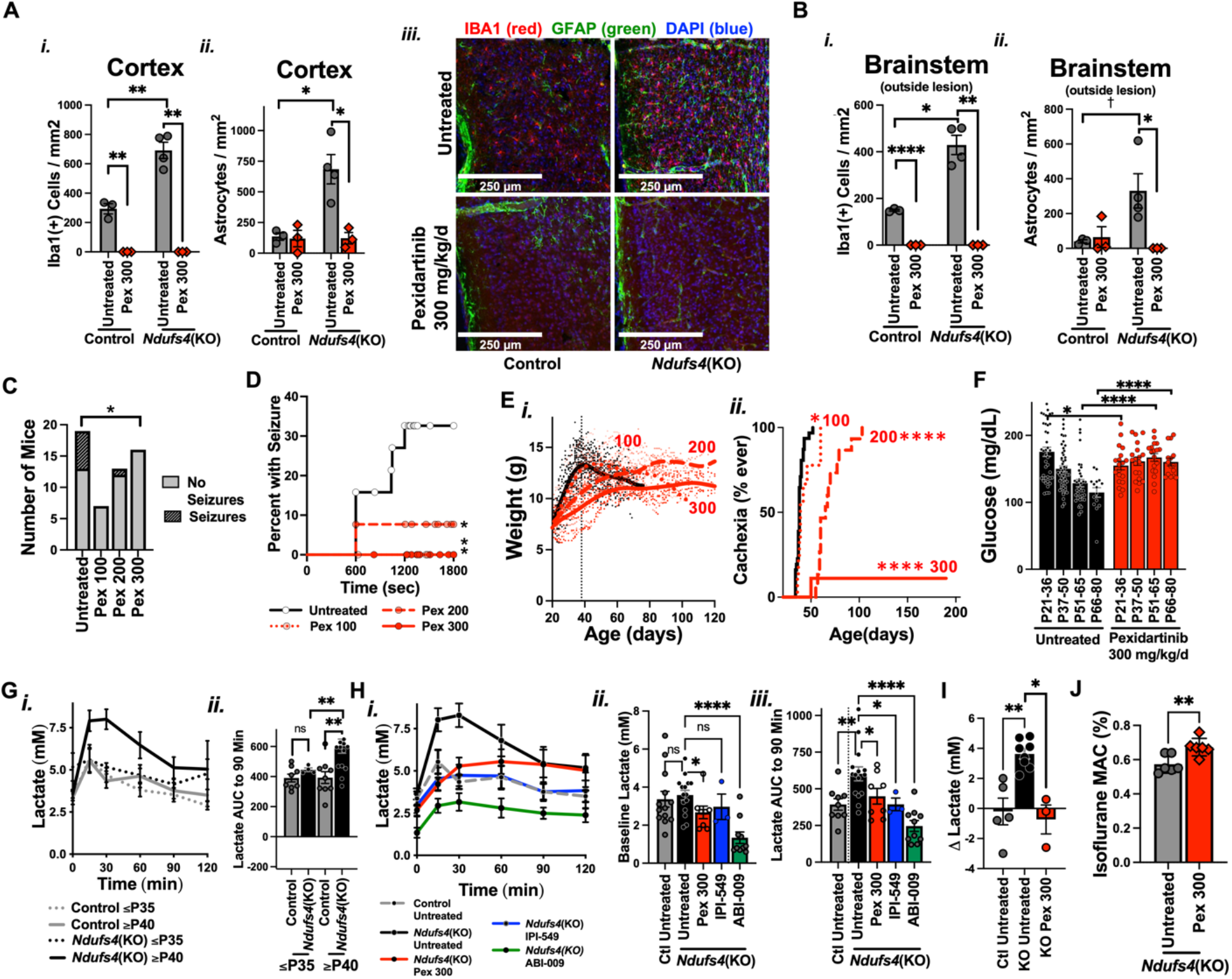
– Leukocyte depletion prevents microgliosis and astrocytosis throughout the brain and rescues a range of systemic symptoms associated with LS in the *Ndufs4*(KO) mice. (A) Quantification of Iba1(+) leukocytes (i) and GFAP(+) astrocytes (ii) in the cortex of control and 300 mg/kg/d pexidartinib treated control and *Ndufs4*(KO) (see ***Methods***). (iii) Representative images of cortex. Datapoints represent individual animals, error bars show SEM. *p<0.05 and **p<0.005 by unpaired, unequal variances (Welch’s), t-test. (B) Quantification of Iba1(+) leukocytes (i) and GFAP(+) astrocytes (ii) in brainstem regions outside of overt lesions in control and 300 mg/kg/d pexidartinib treated control and *Ndufs4*(KO). See ***Fig. 2*** for representative images. Datapoints represent individual animals, error bars show SEM. *p<0.05 and **p<0.005 by unpaired, unequal variances (Welch’s), t-test. †p<0.05 by student’s t-test, p<0.06 by Welch’s t-test. (C) Frequency of seizures at P30 in the rotarod assay by treatment (*Ndufs4*(KO) genotype only – seizures not observed in control mice). *p<0.05 by Fisher’s exact test. (D) Time to seizure for animals in *Ndufs4*(KO) mice in the rotarod assay at P30. All datapoints shown (none censored). *p<0.05 by log-rank test. (E) Weight and cachexia. (i) and (ii) Black line – control treated, dotted red line – 100 mg/kg/d, dashed red line – 200 mg/kg/d, solid red line 300 mg/kg/d treated *Ndufs4*(KO) animals. (i) Scatter plots of *Ndufs4*(KO) mouse weight as a function of age and treatment, with local regression (Lowess) curve overlayed to display population trends. (ii) Cachexia onset (see ***Fig. 1*** and ***Methods***) in control and pexidartinib treated *Ndufs4*(KO) mice. *p<0.05, ****p<0.00005 by Log-rank test. (F) Blood glucose by age in control and pexidartinib treated *Ndufs4*(KO) animals. Each point represents the median value measured for one animal during the given time period (datapoints are biological replicates). *p<0.05 and ****p<0.0001 by unpaired, unequal variances (Welch’s) t-test. (G) Blood lactate levels in response to a glucose bolus (2 g/kg) in control and *Ndufs4*(KO) mice at pre-disease (P25) and early disease (P45) ages. (i) Time-course of lactate levels and (ii) total area under the curve (AUC) for blood lactate from 0-90 min. Error bars represent SEM, **p<0.005 by unpaired, unequal variances (Welch’s) t-test. (H) Blood lactate levels in response to a glucose bolus (2 g/kg) in untreated control and *Ndufs4*(KO) mice and *Ndufs4*(KO) mice treated with pexidartinib (300 mg/kg/d in chow), IPI-549 (100 mg/kg/d in chow), or rapamycin (ABI-009 formulation, 8 mg/kg/d by IP injection). (i) Time-course of lactate levels, (ii) baseline lactate, and (iii) total area under the curve (AUC) for blood lactate from 0-90 min. Error bars represent SEM, *p<0.05, **p<0.005, ****p<0.0001 by unpaired, unequal variances (Welch’s) t-test. (I) Change in blood lactate concentration in control and *Ndufs4*(KO) mice after a 30 min exposure to 0.4% isoflurane and the impact of treatment with 300 mg/kg/d pexidartinib. *p<0.05, **p<0.005 by unpaired, unequal variances (Welch’s) t-test. (J) Mean alveolar anesthetic concentration (MAC) of isoflurane associated with anesthesia in control and 300 mg/kg/d pexidartinib treated *Ndufs4*(KO) mice. **p<0.005 by unpaired, unequal variances (Welch’s) t-test.

### Pexidartinib treatment prevents rotarod induced seizures

Epileptic seizures are common in LS and often refractory to standard therapies (*19*). Given presence of inflammation responsive to pexidartinib in multiple CNS regions, we next wondered if LS features not previously linked to the lesions, such as seizures, might also be rescued by targeting leukocytes.

The rotarod assay provides a mild epileptogenic stimulus in the *Ndufs4*(KO): seizures occur in ∼30% of untreated *Ndufs4*(KO) mice, but not observed in control animals, during rotarod at age P30 (***Fig. 3C***, see ***Methods***). To determine whether pexidartinib attenuates seizures, we performed rotorod on control and pexidartinib treated *Ndufs4*(KO) mice. Incidence of rotorod induced seizures, and time-to-seizure, were both significantly reduced by pexidartinib (***Fig. 3C-D***).

### Pexidartinib treatment rescues hypoglycemia and cachexia, and hyperlactemia is prevented by rapamycin, IPI-549, and pexidartinib

Metabolic features are major sequelae of MD. We next considered the possibility that leukocyte activity may drive some of the metabolic sequelae of LS.

As noted, treatment with pexidartinib prevented both cachexia and hypoglycemia in the *Ndufs4*(KO) mice in a dose-dependent manner (***Fig. 3E-F***) (*5*), consistent with leukocytes playing a role in metabolic derangements.

Abnormally high blood or CNS lactate (often by lactate/pyruvate ratio) is frequently reported in LS and some other forms of MD (*20-23*). In addition, increased lactate is a feature of LS CNS lesions when imaged by magnetic resonance spectroscopy (MRS), and increased intracerebral lactate by MRS has been reported in *Ndufs4*(KO) mice (*24-26*). The *cellular* origins of increased lactate in MD have not been defined. Leukocytes are highly glycolytic (*27*), so a role for leukocytes in contributing to increases in lactate appears reasonable.

To probe hyperlactemia in untreated *Ndufs4*(KO) mice, we measured blood lactate levels at baseline and in response to a glucose bolus in a glucose tolerance test (GTT) paradigm. Clearance of glucose is not significantly altered in the *Ndufs4*(KO), and blood lactate is not increased in *Ndufs4*(KO) animals at baseline, but exposure of *Ndufs4*(KO) animals to a glucose bolus resulted in a significant rise in blood lactate not observed in control mice (***Fig. 3G-H, Fig. S7***). This glucose-induced hyperlactemia also occurred only in *Ndufs4*(KO) mice older than P35, the approximate age of CNS symptom onset. Strikingly, pexidartinib treatment prevented the glucose-induced hyperlactemia.

Given the clinical relevance of hyperlactemia and the novelty of this finding, we sought to further probe the relationship between the mTOR, PI3Kγ, and leukocytes in mediating hyperlactemia in response to glucose. We next tested animals treated with ABI-009 or IPI-549, finding that both compounds prevented hyperlactemia in response to a glucose bolus (***Fig. 3H***).

### *Pexidartinib prevents volatile anesthetic induced hyperlactemia and anesthesia hypersensitivity in the* Ndufs4*(KO)* mice

Hypersensitivity to VAs is a feature of some forms of MD, and no intervention has yet been shown to attenuate MD VA hypersensitivity (*28*). Hypersensitivity to, and toxicity from, VAs is conserved from invertebrates to mammals, including *Ndufs4*(KO) mice and human LS patients with ETC CI defects (*28, 29*). As part of an unrelated anesthesia study, we recently found that low-dose isoflurane exposure leads to a blood lactate spike in *Ndufs4*(KO) mice compared to controls (***Fig. 3I***). Given the impact of pexidartinib on lactate production during the GTT, we tested whether pexidartinib might impact isoflurane induced hyperlactemia. Remarkably, treatment fully suppressed this lactate spike in *Ndufs4*(KO) animals (***Fig. 3I***).

To test whether leukocyte depletion might impact hypersensitivity to sedation itself, we measured the minimum alveolar anesthetic concentration (MAC) (see (*30*)) of isoflurane in control and 300 mg/kg/day pexidartinib treated *Ndufs4*(KO) animals (see ***Methods*** for details). We focused on P30 to assess sensitivity without overt neurologic disease. Pexidartinib provided a modest, but statistically significant, attenuation of the hypersensitivity in the *Ndufs4*(KO) mice (***Fig. 3E***).

### *Pexidartinib significantly increases* Ndufs4(KO) *lifespan, while drug toxicity limits survival in pexidartinib treated animals*

In addition to attenuating multiple features of disease, pexidartinib substantially extended survival of *Ndufs4*(KO) animals in dose-dependent manner from ∼100-300 mg/kg/day (***Fig. 4A***). Critically, the survival curves of control and *Ndufs4*(KO) animals treated with 300 mg/kg/d pexidartinib overlap (at ∼300% increase in survival among the *Ndufs4*(KO) animals), indicating that the drug, rather than underlying MD, is limiting lifespan in this context. Consistent with this notion, the majority of high dose pexidartinib animals did not show overt signs of severe neurologic sequelae prior to death; the proximal cause of death was unclear for both *Ndufs4*(KO) and control animals, with animals dying spontaneously without severe disease (see ***Discussion***).

**Figure 4.**
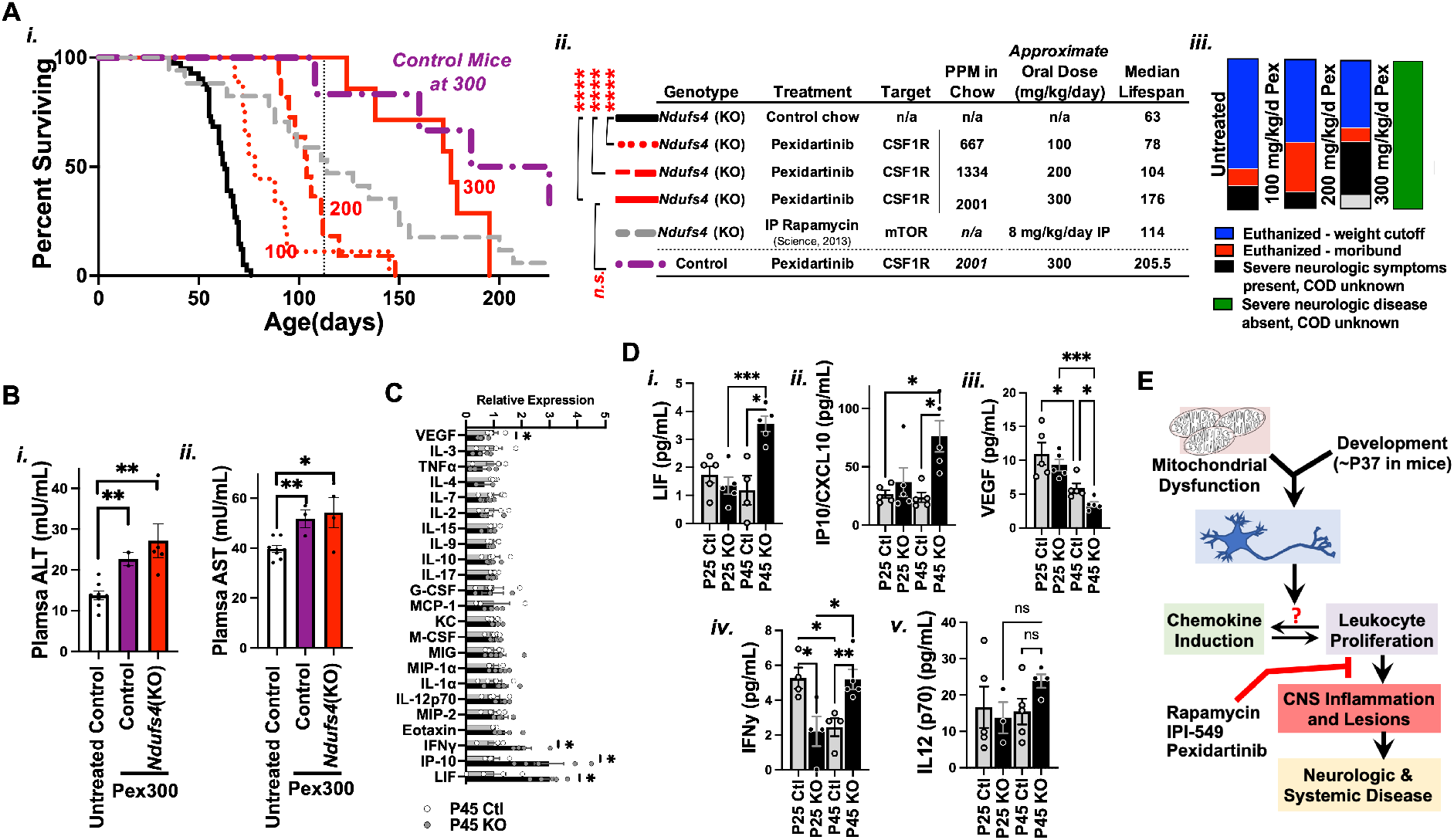
– Pexidartinib dose-dependently extends *Ndufs4*(KO) survival, with survival limited by drug toxicity rather than CNS disease. (A) Survival and cause of death in *Ndufs4*(KO) mice treated with increasing doses of pexidartinib. (i) Survival curves. Black line – control treated *Ndufs4*(KO). Red dotted, dashed, and solid lines - *Ndufs4*(KO) mice treated with 100, 200, or 300 mg/kg/d pexidartinib, respectively. Purple dashed/dotted line – control animals treated with 300 mg/kg/d pexidartinib. Grey dashed line – rapamycin lifespan (for reference). (ii) Median lifespans and dosing data associated with (i). (iii) Cause of death for *Ndufs4*(KO) animals in control and pexidartinib treatment groups (all control animals on pexidartinib 300 mg/kg/d diet of unknown causes with no evident illness). (B) Plasma ALT (i) and AST (ii) levels as determined by enzymatic activity assay. *p<0.05 and **p<0.005 by unpaired, unequal variances (Welch’s) t-test. (C) Levels of 23 chemokines (see ***Methods***) in brainstem of P45 *Ndufs4*(KO) and control animals. *p≤0.05 by unpaired, unequal variances (Welch’s) t-test. (D) Concentrations of select chemokines in brainstem of control and *Ndufs4*(KO) animals at P25 and P45. (i) Leukemia inhibitory factor (LIF), (ii) Interferon gamma-induced protein 10 (IP-10), (iii) Vascular endothelial growth factor (VEGF), and (iv) interferon *γ* (IFN *γ*). *p<0.05, **p<0.005, and ***p<0.0005 by unpaired, unequal variances (Welch’s) t-test. (E) Our data, taken with recent findings linking disease to glutamatergic neurons, lead to a new model for the pathogenesis of disease in LS. In this model, CNS lesions, and at least a portion of other systemic sequelae, are not the direct result of energetic stress or ROS, but instead are causally downstream of immune involvement. Key remaining questions include 1) what specific aspect of mitochondrial dysfunction leads to inflammatory signaling? 2) what is the exact relationship between chemokine production and leukocyte proliferation? and 3) what defines the specific age of onset (∼P37) in the *Ndufs4*(KO) modelã

Chronic administration with pexidartinib is known to cause liver damage, and is associated with risk of serious cholestatic or mixed liver injury, and hepatic function is carefully monitored in patients taking pexidartinib (*31*). Accordingly, we tested blood alanine aminotransferase (ALT) and aspartate aminotransferase (AST), markers of hepatic damage elevated in pexidartinib treated human patients (*32, 33*), in control and *Ndufs4*(KO) mice treated with 300 mg/kg/day pexidartinib to assess whether hepatotoxicity may contribute to early mortality. Consistent with this notion, pexidartinib treated control and *Ndufs4*(KO) mice at ∼P150 have significantly elevated ALT and AST compared to age-matched untreated control animals (note: *Ndufs4*(KO) mice universally perish prior to this age, precluding inclusion in this dataset) (***Fig. 4B***) (see ***Discussion***).

### *Inflammatory chemokines are significantly increased in* Ndufs4(KO) *brainstem only at ages associated with disease*

Taken together, the rapamycin, IPI-549, and pexidartinib data reveal that leukocytes proliferation is a key causal step in the pathogenesis of LS. Furthermore, the benefits of IPI-549 specifically implicate extracellular signaling, suggesting that specific chemokine factor(s) may be involved. If so, the specific age of onset of disease in the *Ndufs4*(KO) mice, and postnatal onset of LS in humans, might suggest one or more inflammatory factors are induced at a post-natal age, driving disease.

To probe these possibilities, we performed targeted cytokine profiling of control and *Ndufs4*(KO) brainstem from P25 (prior to overt disease) and P45 (early in disease progression) animals (see ***Fig. 1A***). Among the 23 factors in our panel reliably detected above background, four showed significantly altered expression in *Ndufs4*(KO) mice compared to controls at P45: interferon gamma (IFNγ), IFNγ-Induced Protein 10 (IP-10/CXCL10), and Leukemia Inhibitory Factor (LIF) were all increased in the *Ndufs4*(KO) animals at P45, while vascular endothelial growth factor (VEGF) was significantly reduced. IL-12 (p70) was also increased in *Ndufs4*(KO) mice only at P45, though not reaching statistical significance due to high variance (see ***Discussion***).

Notably, CSF-1 itself was not elevated in the *Ndufs4*(KO). CSF1R inhibition impairs leukocyte proliferation, but this strategy is untargeted, impacting all leukocytes. A more targeted therapeutic based on a precise inflammatory factor, if identified, may provide greater benefits with fewer off-target effects.

## Discussion

### A causal role for leukocyte proliferation in the pathobiology of Leigh syndrome

The most important finding in this study is that leukocytes proliferation is a key causal mechanistic step in the pathogenesis of LS. Targeting leukocyte proliferation prevented microgliosis, rescued astrocytosis, led to a complete (as far as we could detect) prevention of CNS lesions, and rescued CNS inflammation outside of overt lesions. Leukocyte depletion prevented sequelae associated with the overt CNS lesions including respiratory dysfunction, balance and movement, and survival. Moreover, leukocyte depletion rescued symptoms of LS not directly attributed to the CNS lesions – hypoglycemia, cachexia, hyperlactemia in response to a glucose bolus, hyperlactemia in response to anesthesia, seizures, and anesthesia sensitivity. These findings dramatically shift our understanding of LS.

### Central vs peripheral leukocytes and microglia vs peripheral macrophages

The relative contribution of central versus peripheral leukocytes to individual disease features remains to be determined, and will require significant further study.

The precise nature of immune cells in neuroinflammation, including the CNS lesions, is also not yet known. While the overt lesions have been described as involving ‘microgliosis,’ it is critical to note that the first study in 1951 by Denis Leigh described microgliosis based on only Nissl staining, which is wholly non-specific (*34*). To our knowledge, all studies utilizing the *Ndufs4*(KO) have relied on Iba1 staining, which was utilized by the lab which generated the model (*35*). Iba1 is, unfortunately, also not microglia-specific - Iba1 is expressed in all macrophages, has recently been termed a pan-macrophage marker, and is expressed in other cell types, including antigen presenting cells, under certain conditions (*36, 37*). Unfortunately, no simple immunohistochemical approach alone can determine the peripheral versus central immune composition of CNS lesions in this model (TMEM119, for example, is microglia specific, but is downregulated in inflammatory microglia, so Iba1(+)/TMEM119(-) staining does not rule out microglia (*38*)). Significant future efforts will need to address this outstanding question.

While microglia are a likely suspect in driving at least some CNS pathologies, we are careful to present our findings in terms of leukocytes broadly – as with Iba1, rapamycin, IPI-549, and pexidartinib are not specific to microglia, and our data alone cannot distinguish between resident microglia and peripheral leukocytes.

### A mechanism for mTOR inhibition in Leigh syndrome

Determining the mechanisms underlying the benefits of rapamycin in the *Ndufs4*(KO) model has been an active area of research since the effects of the drug were published in 2013 (*4-6, 39-44*). The data presented here reveal that the benefits of mTOR inhibition in the *Ndufs4*(KO) result *primarily* from immune modulation. However, it is important to note that mTOR inhibitors have shown beneficial effects in a variety of MD models including cultured cells and invertebrates (*6, 40-44*). mTOR inhibition is highly pleiotropic, and it is clear that other mTOR regulated processes, such as metabolism and autophagy, mediate the benefits of targeting mTOR in other settings. A full review of the pleiotropic effects of mTOR inhibition and the models that benefit from targeting mTOR is not possible here, but it is clear that the immune mechanism we identify is not shared in all settings of mitochondrial dysfunction. On the other hand, immune activation may play a previously unappreciated role in the presentation of some other forms of MD, a possibility warranting further attention.

Prior work aimed at defining the role of mTOR in the *Ndufs4* (KO) has led to the identification of multiple putative pathways mediating the benefits of mTOR inhibition. These include regulation of metabolism, neurotransmitter abundance, PCK activity, and regulation of transcription (*5, 6, 45-48*). Without attempting to review this literature in depth we note two important observations: 1) none of these targets matched or surpassed the benefits of mTOR inhibition; 2) none are inconsistent with the finding we report here. In the PKC study, for example, the authors note the role of PKC downstream of mTOR in innate and adaptive immune regulation, and suggest that modulation of immune function may mediate the effects. We believe our data provide a comprehensive model accounting for most or all of this literature.

### Relation to other targets

In other studies in the *Ndufs4* (KO) model, rescue of NADH redox by yeast Ndi1 expression has been shown to rescue lifespan but not ataxia (*49*), while chronic mild hypoxia provides substantial benefits to disease (*50, 51*). Ndi1 expression is presumed to act on primary molecular sequela (redox and redox regulated processes), preventing downstream cell- and tissue-pathologies. Chronic hypoxia may act upstream of immune involvement by preventing primary molecular sequelae (perhaps by preventing the ‘oxygen toxicity’ suggested by the authors of these studies), but might also act robustly on the immune response. The relationship between NADH redox, hypoxia, and the mechanisms we report here.

### A role for leukocytes in blood lactate

The rescue of hyperlactemia induced by glucose or isoflurane was perhaps the most surprising benefit of pexidartinib. These data suggest that leukocytes, or the consequences of tissue inflammation they drive (e.g. tissue ischemia), may be a major source of blood lactate. Alternatively, degeneration of CNS regions or peripheral tissues involved in whole-body metabolism may impact the metabolic fate of glucose. These findings may be relevant to other forms of MD where hyperlactemia is present.

### A new model for the pathogenesis of Leigh syndrome

Onset of symptoms in the *Ndufs4*(KO) occurs at ∼P37 (see ***Fig. 1***), reminiscent of human LS, where patients are often overtly healthy at birth. Here, we show that pro-inflammatory chemokines are increased in mice after this age of symptom onset. Prior studies have demonstrated that neuron specific (Nestin-Cre) or glutamatergic neuron specific (VGlut2-Cre) deletion of *Ndufs4* results in a near-complete recapitulation of the disease including CNS lesions, cachexia, metabolic dysfunction, behavioral deficits, and reduced survival, while *Ndufs4* deletion in GABAergic or cholinergic neurons does not (*35, 45, 52*). Considering these data in the context of this study, we can assemble a model for the pathogenesis of disease in the *Ndufs4*(KO): the presence of mitochondrial dysfunction during post-natal development causes cellular sequelae in VGlut2 positive neurons (the nature of which is yet to be defined), resulting in the induction of IFNγ and related chemokines. These chemokines drive leukocyte proliferation, which cause tissue degeneration, astrocytosis, CNS dysfunction, and systemic symptoms (***Fig. 4***).

### New insights into mysteries surrounding LS

By revealing a necessary step in the pathobiology of LS other than ETC function, energetics, or ROS, our data may resolve key mysteries regarding this disease. In particular: 1) why is there a general sparing of some tissues with high energy requirements, as in the lack of any overt cardiac phenotype in cardiac specific *Ndufs4*(KO) in the absence of a stressor (*53-55*); 2) why does LS often develop postnatally with no symptoms at birth? The enigmatic site specificity of lesions in LS may be elucidated when the initiating signal is identified, but significant additional work will be necessary to probe this lead.

Viral infection or fever has been reported to often occur just prior to symptom onset in MDs, and in certain other forms of acute focal necrotizing encephalopathy in children (*56-62*). This link in MD has been attributed, by some, to energetic stress caused by mobilizing an immune response (*58*). Though speculative, our data may suggest that induction of pro-inflammatory signaling contributes to this observed link.

### Therapeutic implications

Our findings may have significant therapeutic implications. Clinical translation of basic research is exceptionally difficult in rare diseases (*63-66*), compounded in LS by post-natal onset, complications of diagnosis, and the clinical and genetic heterogeneity of the disorder (over 75 causal genes have been identified) (*67-69*). mTOR inhibition was the first small molecule shown to attenuate disease in the *Ndufs4*(KO), and has since been found to benefit at least some patients with MD (*6, 7, 40*). However, the translational potential of mTOR inhibition is limited by the incomplete rescue in the animal model, the lack of a mechanistic understanding of the beneficial effects, and the well-documented off-target effects of mTOR inhibitors.

While we do not suggest that the specific compounds used here will, or should, be trialed in patients, the discovery that leukocyte proliferation mediates disease opens the door for an entirely new pathway for therapeutic targets. A number of well-characterized clinically approved agents already exist for targeting immune function. Moreover, follow-up studies have a high probability of identifying of novel targets in the observed immune activation with greater potency and reduced off-target effects.

We suspect that leukocyte proliferation contributes to the pathogenesis of other MDs involving CNS lesions, and may extend to MD syndromes where inflammation is present. Determining the generalizability of our findings to genetically distinct forms of LS and other MDs will require experimental work and will be an important aspect of future studies.

### Inflammatory clues

Our data identify a potent and effective new pathway for therapeutic targeting. The benefits of pexidartinib in our preclinical model appear limited by toxicity. Improved CNS targeting to lower necessary dosing, identification of more specific targets for intervention (such as inhibitors of CXCL3, the receptor for IP10), testing of combinatorial therapies, and chemical modifications to current inhibitors to lower toxicity may lead to robust small molecule therapeutic options. By identifying a specific biological process with many druggable regulatory players we hope to have opened the floodgates for innovation into novel treatments for this disease.

Our targeted chemokine panel data provide only a peek into the inflammatory mechanisms at play, and further exploration of inflammatory signaling networks beyond the scope of this current manuscript, but we consider these data of great potential value for future studies. In particular, the known relationship between the identified factors of interest suggests a discrete mechanism for immune activation. IL-12 produced by activated antigen-presenting cells drives expression of IFNγ and inhibits VEGF expression, while IFNγ induces both IP-10 and LIF (*70-72*). Moreover, this IL-12 chemokine axis plays a prominent role in graft-versus-host (GVH) disease (*73, 74*); prevention of GVHD is, of course, a clinically approved use of mTOR inhibitors (*75*).

As mentioned, it is particularly notable that human disease often first appears after a viral infection or fever. IL-12, IFNγ, and IP-10 suggest that innate immune responses provide a mechanistic link between infection and disease onset, though why IFNγ and related factors are upregulated at a specific age in mice, and whether these same factors are in fact increased in human LS patients, remains to be determined. Given the role of IL-12/IFNγ/IP10 in responding to pathogens, it seems reasonable to hypothesize that mitochondrial defects lead to the aberrant sensing of some dysfunctional mitochondrial component (perhaps free mtDNA or misfolded mitochondrial proteins) as ‘foreign’ by the immune machinery.

The recently described cGAS/STING (cyclic GMP-AMP synthase (cGAS)/stimulator of interferon genes) pathway, which senses intracellular DNA and induces IFN responses, is a top candidate for further study; however, cGAS/STING drives type I interferon production, and has only been shown to impact IFNγ through indirect mechanisms (i.e. regulation of IFNγ producing T cells), so the relationship is unlikely to be direct (*76, 77*).

### Caveats of pharmacologic studies

The positive findings with p110γ and CSF1R inhibitors, not the negative results with the p110α, p110β, and p110δ inhibitors, underlie the significance of this study. While all PI3K isoform inhibitors are of similarly potency and tissue accumulation, and our data suggest p110α, p110β, and p110δ are not effective targets, we cannot rule out the possibility that alternative targeting methods might be more effective. Conversely, though these inhibitors are highly specific, the modest benefits of p110α, p110β, and p110δ inhibition may simply be the result of off-target effects on p110γ.

### Unanswered questions

While we believe our data represent a significant step forward in our understanding of LS, and perhaps MD more broadly, key questions remain. What *precise* cells initiate CNS inflammation? What underlies immune signaling induction, and how does altered mitochondrial function drive this signal? Is inflammation restricted to the CNS, or are some non-CNS symptoms the result of peripheral inflammation? Does inflammation play a role in other forms of MD? Finally, what link, if any, is there between inflammation and chronic hypoxia, the most potent therapeutic strategy yet identified in the *Ndufs4* (KO) (*51*)? The changes we observe in VEGF suggest there may be a link, but the nature of the relationship between our findings and hypoxia remains to be determined. Answers to these questions will shed new light on the interplay between mitochondria, the immune system, and systemic disease.

## Materials and Methods

### Ethical statement, animal use, breeding, care, and euthanasia criteria for survival studies

All care of experimental animals was in accordance with the Seattle Children’s Research Institute guidelines and experiments were performed as approved by the Institutional Animal Care and Use Committee.

*Ndufs4*(+/-) mice were bred to produce *Ndfus4*(KO) (*Ndufs4*(-/-)) offspring. Mice were weaned at 20-21 days of age. *Ndufs4*(KO) animals were housed with a minimum of one control littermate for warmth and stimulation. Mice were weighed, and health assessed, a minimum of 3 times per week, every other day (daily for IP injected mice, see below). Where longitudinal blood point-of-care data was collected, this was performed during these health checks. Wetted chow in a dish on the bottom of each cage, and in-cage water bottles, were provided to all cages housing *Ndufs4*(KO) mice following the onset of disease so that the ability to find or reach food or water was not a limiting factor for survival.

Mice were euthanized if they showed a 20% loss in maximum body weight, immobility, or were found prostrate or unconscious.

As we and others have previously reported, *Ndufs4* deletion is a recessive defect, and heterozygosity results in no reported or observed phenotypes, including no detectable defects in ETC CI activity, so controls for this dataset include both heterozygous and wildtype mice.

All experimental mice were fed PicoLab Diet 5058; pharmacologic agents were compounded into this diet (see below).

### Longitudinal assessments of neurological disease symptoms

Clasping, ataxia, and circling were assessed by visual scoring, as previously described. As disease progresses in the *Ndufs4*(KO) mice animals display intermittent/transient improvement of symptoms. For these studies, we simplified our analysis to simply report whether animals had ever reported the symptom – requiring that the symptom was observed for two or more consecutive days to minimize spurious reporting. Transient loss of symptoms was not included in these data. For observational assessments, lab-wide quality control discussions occurred frequently to ensure consistency between technicians/researchers. Technicians contributed equally to each treatment group to minimize any potential bias between individuals.

Cachexia in ***Fig. 1*** and ***Fig. 3*** is the day of life when weight peaks prior to the progressive weight loss which occurs in untreated *Ndufs4*(KO) animals.

### Replicate numbers and control group

Animal numbers for each dataset are noted in figure legends, and whenever possible all individual datapoints are shown.

Animals were added to the control treatment groups across the duration of the experiments presented here to ensure that no shifts in colony survival, behavior, etc, occurred during the course of these studies. Accordingly, control treatment groups generally contain larger cohorts than the individual treatments.

### Reagents

Laboratory chemicals were purchased from Sigma. Pexidartinib, IPI-549, CAL-101, and GSK2636771 were purchased from MedKoo Biosciences (cat. #’s 206178, 206618, 200586, and 205844, respectively). BYL719 was purchased from AChemBlock (cat. #R16000).

Doses of each drug was based on published data for use in animals. Tissue concentrations reached biologically relevant levels for each of the tested inhibitors, see ***Figs. S2, S5***.

### Pharmacologic interventions and chow formulation

Standard mouse chow was ground to a power and mixed with drugs at the doses desired (see inset table in ***Fig. 1*** for PPM equivalents). 300 mL of 1% agar melted in sterile water was added per kilogram of powdered chow and the mixture was pelleted and baked at 37°C for 3-5 hours to dry. Pellets were then stored at 4°C for short term (up to 90 days) or −20°C for long-term (up to 6 months) storage. We have previously shown that this processing has no impact on animal health or survival (*4*).

Daily food consumption values were estimated based on data in ***Fig. S1*** – we used the value of (0.15 g food / g mouse / day), at the lower end of the daily food consumption estimate range, to ensure mice received the desired level of drug.

### ABI-009

Rapamycin was provided in the form of ABI-009 (nab-rapamycin), an albumin encapsulated water-soluble formulation. ABI-009 was provided as a gift by Aadi, LLC, in lyophilized form. ABI-009 was resuspended to 1.2 mg/mL rapamycin in 1X PBS. This solution was sterile filtered and stored in aliquots at −80 degrees Celsius. ABI-009 was administered at 66 µL per 10 g for a final dose of 8 mg/kg/day rapamycin, as in our prior studies (*4, 5*).

### Point-of-care glucose and lactate testing

Blood glucose and lactate measurements were collected using point-of-care meters (Prodigy Autocode glucose meter, product #51850-3466188; Nova Biomedical lactate assay meter, product #40828) and the tail-prick method, as previously described (*78*).

### Rotarod and rotarod seizures

Rotarod assays were performed using a Med Associates Inc ENV-571M single lane rotarod with a black mat installed in the floor of the lane to reduce visibility of the bottom. Assays were performed by placing animals onto an already rotating rod and timing latency to fall, with a steady rotation speed set to 6 rpm (controlled by attached laptop and Med Associates software). The maximum time of each trial was 10 min, with the trial ending at that time if mice were still on the rotarod. For each assay, three trials were performed, with a minimum of 5 min between each assay. The best of the three trials was reported for the rotarod data. As detailed in the text and figure legends, rotarod was performed each 10 days of age from P30; mice in a given age group are the reported age +/- 1 day.

Mice were monitored throughout each assay for seizure activity, and rotarod performance trials were ended if a seizure was observed. Mice were considered to be showing seizure activity if any symptoms on the Racine behavioral scale or Pinel and Rovner scale were observed: abnormal oroalimentary movements (dropping of the jaw repeatedly, atypical gnawing or chewing movements), repeat head nodding, anterior limb clonus (twitching/jumping while making no contact with the face), dorsal extension/rearing, loss of balance and falling (observed when mouse was off the rotarod), violently running/jumping. Critically, no attempt was made to assess seizures on the seizure scales for these studies – only presence or absence was noted.

While blinding was often impossible (BYL719 treatment leads to extremely small body size, pexidartinib turns fur white, and overtly diseased *Ndufs4*(KO) mice are easily identified), the experiments were scored by technicians with no expectations regarding the outcomes. Technicians trained together and cross-compared observed symptoms to ensure consistent scoring, and analysis of each treatment group was roughly evenly split among technicians involved in these studies to prevent any unaccounted-for observer bias from impacting any individual treatment group.

### Glucose tolerance test and glucose induced lactate test

Prior to performing the GTT, mice were fasted for 4 hours, from 10 AM-2 PM. The fasting duration was limited to 4 hours given the sensitivity of untreated *Ndufs4*(KO) animals to hypoglycemia and related sequelae (dysregulation of body temperature, hypoglycemia induced seizure, etc). For the ABI-009 cohort, mice received their daily ABI-009 treatment at the start of this fasting period. At the end of the 4 hour fast, baseline blood glucose and lactate values were collecting using point-of-care meters and the tail-prick method. Mice were then injected by IP with 2 g/kg dextrose (10 microliters/gram body weight of 200mg/mL glucose in 1x PBS, 0.2 micron sterile filtered) using an insulin syringe (31-gauge, 6 mm length, 3/10 cc, BD Veo Ultra-Fine needle). Glucose and lactate levels were measured using a minimally invasive tail-prick method at designated timepoints post-injection.

### Isoflurane induced hyperlactemia assays

All of these experiments were performed at P50. Mice were subject to normal daily monitoring (see *survival studies*) up until P50. At P50, baseline blood glucose, lactate, and β-HB levels were measured using point-of-care meters using the tail-prick method, as in detailed. Mice were then immediately exposed to 0.4% isoflurane or carrier gas only (100% oxygen) for 30 min. At the 30 minutes, blood glucose, lactate, and β-HB were measured again.

### Anesthetic sensitivity by minimum alveolar anesthetic concentration

The minimum alveolar anesthetic concentration (MAC) was determined as previously described (Sonner et al. 1999 and 2000). In brief, P30 mice were placed in an individual gas-tight plastic chamber connected to an oxygen source and carbon dioxide absorber. Isoflurane was provided through a vaporizer (Summit Anesthesia) at a flowrate of 1-2 liters/min and humidified in-line. Mice were kept warm with a circulating water heating pad maintained at 38°C (Adroit Medical, HTP-1500). Anesthetic concentration was adjusted in steps of 0.2%, equilibrated for 10 minutes, and was monitored with an in-line analyzer (AA-8000 Anesthetic Agent Analyzer, BC Biomedical). MAC was calculated as the mean anesthetic concentration bracketing the animal’s response and non-response to a nondamaging tail clamp.

### Respiratory measurements

Breathing parameters were recorded from alert, unrestrained, adult (∼P60) mice using whole-body plethysmography. Paired 300 ml recording and reference chambers were continuously ventilated (150 ml/min) with either normal air (79% nitrogen and 21% oxygen) or a hypercapnic gas mixture (74% nitrogen, 21% oxygen, and 5% CO_2_). Pressure differences between the recording and reference chambers were measured (Buxco) and digitized (Axon Instruments) to visualize respiratory pattern, and simultaneous video recordings were performed to differentiate resting breathing activity from exploratory sniffing and grooming behaviors. Untreated and treated (pexidartinib 300 mg/kg/day) control and *Ndufs4*(KO) mice were allowed to acclimate to the chambers for 30-40 min prior to acquisition of 35 min of respiratory activity in normal air. The respiratory response to hypercapnia was then tested by ventilating the chambers with hypercapnic gas for 15 min. Respiratory frequency, and breath-to-breath irregularity scores 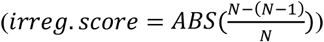 of frequency and amplitude (peak inspiratory airflow) were quantified during periods of resting breathing (pClamp10 software). During hypercapnic challenges, only data during the final 5min of the 15 min period were analyzed. Results were visualized and statistical comparisons between groups were performed in GraphPad Prism9 software.

### Mixed brain cell culturing and staining

Briefly, using sterile technique, brains were removed from neonatal pups into ice-cold HBSS (Gibco cat. #14025076). Brains were washed three times in cold PBS. After the third wash, brains were minced using surgical scissors into small, fine pieces. The minced brain was centrifuged at 100 g for 5 min. Once pelleted, PBS was removed and 0.05% trypsin/EDTA (Gibco) was added. The tissue in trypsin/EDTA was then incubated at 37ºC with gentle shaking for 15-30 min, with intermittent mixing and supplemental mincing, until the solution was homogenous and without tissue segments. After incubation, the minced tissue was placed back into the centrifuge at 100 g 5 min. Pellet was washed 3x with cold PBS before adding fresh media to each tube. The tissue was further dissociated using 1000 µL pipet until the solution became cloudy. The sample was washed through a sterile 70-micron cell strainer (Corning) into a 50 mL falcon tube, then plated onto T75 poly-d-lysine coated Nunclon flasks (Nunc EasyFlask132704) at a ratio of one brain to two flasks. After 24 hours, dead cell/debri were washed and media replaced. Cells were then grown for 1 additional day, split onto multi-well plates with poly-d-lysine coated coverslips (Cellware 12mm round coverslips) at approximately 25% confluence with ABI-009 or pexidartinib (for dose-response) added. Media was replaced (with pharmacologic agents) every 3 days, and cells were fixed in ice-cold 3.7% PFA on day 7.

Fixed cell slides were washed 3X with 1X PBS followed by a 5 min incubation in 0.2% Triton-X100/1XPBS at room temperature. Slides were washed again 3X in 1XPBS, blocked for 30 min at room temperature in 10% rabbit serum, and then incubated in antibody solution overnight at 4ºC protected from light. Antibody solution consisted of blocking solution with rabbit anti-Iba1-635 (Wako, cat. # 013-26471) at 1:400 and DAPI (Sigma, cat. # D9542) at 10 µg/mL.

Cell media for these experiments consisted of 500 mL of DMEM (Gibco ref#11995-065), 56.2 mL of One-Shot Fetal Bovine Serum (FBS) (Gibco cat. #16000-077), and 5.62 mL of penicillin/streptomycin, 10,000 U/mL (Gibco cat. # 15140122).

Pharmacologic agents were sourced as listed below, dissolved to 1000X in DMSO, and added at 1:1000 to media for working media. DMSO was added to the ‘no drug’ wells. These experiments were repeated 3 times with similar results. Data in ***Fig. 2*** are derived from biological replicates from the same experiment.

### Brain immunological staining and microscopy

Brains were fixed for 48 hours in 10% formalin at 4º C. Following fixation, brains were moved to a cryoprotectant solution (30% sucrose, 1% DMSO, 100 μM glycine, 1X PBS, 0.45 μm filtered, pH 7.5), and stored for over 48 hours, until the fixed tissues sank to the bottom. Tissues were then placed in OCT media (Tissue-Tek OCT compound, Sakura 0004348–01), frozen in cryoblock holders on dry ice, and stored at −80 until sectioned for staining. Cryoblocks were cut at 50 μm thickness using a Leica CM30505 cryostat set at −40 degrees C. Slices were moved to 1X PBS and stored at 4 degrees C until used for staining. Prior to staining, slices were mounted on slides and briefly dried to adhere.

Antibody staining was performed as follows: slides were first incubated in an antigen retrieval and permeabilization buffer (0.05% Triton X-100, 50 μM digitonin, 10 mM Tris-HCl, 1 mM EDTA, pH 9.0) at 60ºC overnight in a white-light LED illuminated box (to promote photobleaching of tissue autofluorescence). To reduce formaldehyde induced background fluorescence, slides were treated with sodium borohydride in ice cold PBS, added at 1 mg/mL immediately before incubation, on ice for 30 min, then moved to 10 mM glycine 1XPBS, pH 7.4, for 5 min at room temperature. Lipid background fluorescence was then blocked by incubating slides in 0.2 um filtered Sudan Black B solution (5 mg/mL in 70% ethanol) overnight at room temperature. Slides were then rinsed twice, 5 min each, in 1X PBS. Excess fluid was wiped from the slide, and the tissue was circled using Liquid Blocker PAP pen (Fisher Scientific, NC9827128) to hold staining solutions. Slides were blocked for 15 min at room temperature in 1X PBS with 10% rabbit serum (Gibco, 16120–099) then stained overnight at 4 °C in a mixture of rabbit anti-IBA1-fluorochrome 635 conjugated (WAKO 013-26471) at 1:300, mouse anti-GFAP, Alexa555 conjugated (Cell Signaling, 3656S) at 1:300, and DAPI (Sigma, D9542) at 1 μg/mL. The following day, slides were washed 3X 5 min in 1X PBS then mounted in aqueous mounting media (Fluoromount-G), coverslipped, and stored at 4 °C protected from light until imaging.

Slices were imaged on a Zeiss LSM 710 confocal microscope. Images were collected using a 10X dry objective at 0.6X optical zoom, resulting in images of 1417 × 1417 microns in physical area. Images were collected with 8–16 line averages and a line scan speed of 6–7. Channels were set to an optical thickness of 50nm. DAPI was excited at 405 nm, with emission light collected using a sliding filter with the range setting at 424–503 nm. GFAP-Alexa555 was excited with a 543 nm laser, emissions collected at 548–587 nm. Iba-1-635 was excited at 633 nm and emitted light was collected at 641–661 nm.

For quantification, a region approximately 10% of the total image area, with overall structure and cell composition representative of the given image, was selected and GFAP (+) and Iba1(+) cells were counted. While images were collected with identical settings, brightness and contrast were adjusted as needed to detect cell bodies during counting. Only cell bodies of GFAP (+) cells (astrocytes) and Iba1(+) leukocytes were counted (to the best of our ability), with cell extensions from out-of-plane cells ignored. Images in main text figures have identical brightness/contrast settings, but representative olfactory bulb images in supplemental data were adjusted as needed to highlight stained features (there was high variability in intensity of stains in the olfactory bulb, but this does not preclude the analyses in this study, which are based on cell numbers rather than intensity).

### Plasma ALT and AST measures

Small volumes (5-10 µL) of blood were collected using tail-prick and heparinized microhematocrit tubes (Fisher Scientific cat. # 22-362566), placed on ice in 15 mL sample tubes, then moved (by pipette) into 1.5 mL microcentrifuge tubes and stored at −80ºC until used for ALT and AST enzymatic assays. ALT and AST were quantified using colorimetric enzymatic activity assay kits (Sigma, cat. #’s MAK052 and MAK055, respectively) according to manufacturer recommendations. Absorbance was measured on a NanoDrop 1000 (ThermoScientific).

### Chemokine analyses

Brainstem chemokines were analyzed by Eve Biotechnologies using the Millipore MCYTMAG-70K-PX32 Milliplex MAP Mouse Cytokine/Chemokine Magnetic Bead Panel Multiplex panel. Brainstem samples were collected rapidly after euthanasia, flash frozen in liquid nitrogen, and shipped to Eve Biotechnologies on dry ice. Samples were processed and analyzed according to Millipore manufacturer recommendations.

### Drug tissue level quantification

All tissue drug level analyses were performed by the University of Washington Medicinal Chemistry Mass Spectrometry Center commercial service.

Briefly, control mice were fed drug compounded diets ad libitum until post-natal day 30 (10 days, including any ramp up period – see above) to achieve steady state. Animals were euthanized in the early afternoon, tissues flash frozen, stored at −80ºC, and transported to the medicinal chemistry core on dry ice.

#### Sample Prep for all tissues

Tissue samples were prepared by weighing tissue amount and adding 3ml of 1x PBS buffer and kept on dry ice until homogenized. Samples were then homogenized using an OMNI bead disruptor tissue homogenizer, chilled to −15 °C, shaken at 5.8m/s for 45s allowed to settle for 30s, then repeated. Samples were removed and stored at −80C until extracted for analysis.

#### Analysis of Cal101 & IPI549

Cal101, IPI549 and glyburide (internal standard) were measured using high pressure liquid chromatography-mass spectrometry. In summary, the assays were performed on a Water’s Xevo TQ-XS coupled to a Water’s I-Class Ultra high pressure liquid chromatography system (Waters Corporation, Milford, MA, USA). Analytes were monitored in MRM mode (multiple reaction mode) selectively isolating parent m/z 416.1->176.1, 529.2->161.1 and 494.1->369.1, respectively. Chromatographic separation was achieved using a Water’s Acquity T3 C18, 2.1×100mm, 1.8µ column (Waters Corporation, Milford, MA, USA). Using a gradient consisting of 0.1% formic acid in water (H2O) (A) and 0.1% formic acid (FA) in methanol (MeOH) (B) at a flow rate 0.3ml/min. (can provide gradient if needed).

Sample Prep: 10µl of IS (Glyburide-100ng/ml) was added to 20µl of unknown analytical sample or calibration/qc sample (10µl of stock at various concentrations + 10µl of blank matrix). Samples were then vortexed and then 100µl of MeOH was added. Sample was vortexed again and centrifuged at 16.1rfc for 5mins. 50µl of Supernatant was removed and diluted with 50µl H20 in autosampler vials, 5µl injections were made on the platform.

#### Analysis of Pexidartinib

Pexidartinib and glyburide were measured using high pressure liquid chromatography-mass spectrometry. In summary, the assays were performed on a Water’s Xevo TQ-XS coupled to a Water’s I-Class Ultra high-pressure liquid chromatography system (Waters Corporation, Milford, MA, USA). Analytes were monitored in MRM mode (multiple reaction mode) selectively isolating parent m/z 418.2->258.1, and 494.1->369.1, respectively. Chromatographic separation was achieved using a Water’s Acquity BEH C18, 2.1×50mm, 1.7µ column (Waters Corporation, Milford, MA, USA). Using a gradient consisting of 0.1% formic acid in water (H2O) (A) and 0.1% formic acid (FA) in acetonitrile (ACN) (B) at a flow rate 0.3ml/min. (can provide gradient if needed).

10ul of IS (Glyburide-100ng/ml) was added to 2µl of unknown analytical sample or calibration/qc sample (various concentrations spiked into blank matrix). Samples were then vortexed and 60µl of ACN was added. Sample was vortexed again and centrifuged at 16.1rfc for 5mins. 40µl of supernatant was removed and diluted with 40µl of H2O into a limited volume autosampler vial. 2µl injections were made on the platform.

#### Analysis of GSK2636771 and BYL719

GSK2636771, BYL719 and glyburide were measured using high pressure liquid chromatography-mass spectrometry. In summary, the assays were performed on a Water’s Xevo TQ-XS coupled to a Water’s I-Class Ultra high-pressure liquid chromatography system (Waters Corporation, Milford, MA, USA). Analytes were monitored in MRM mode (multiple reaction mode) selectively isolating parent m/z 434.4->215.2, 442.3->328.2 and 494.1->369.1, respectively. Chromatographic separation was achieved using a Water’s Acquity BEH C18, 2.1×50mm, 1.7µ column (Waters Corporation, Milford, MA, USA). Using a gradient consisting of 0.1% formic acid in water (H2O) (A) and 0.1% formic acid (FA) in methanol (MeOH) (B) at a flow rate 0.3ml/min. (can provide gradient if needed).

For blood samples: 20µl of IS (glyburide-100ng/ml) was added to 20µl of unknown analytical sample or calibration/qc samples (10ul of stock at various concentrations + 10µl of blank matrix). Samples were then vortexed (15s) and then 100µl of ACN was added. Samples were vortexed again (15s) and centrifuged at 16.1rfc for 5 mins. 50µl of supernatant was removed and diluted with 50µl of H2O in limited volume autosampler vials, 5µl injections were made on the platform.

For tissue samples: 20µl of IS (glyburide-100ng/ml) was added to 100µl of unknown analytical sample or calibration/qc samples (50µl of stock at various concentrations + 50µl of blank matrix). Samples were vortexed (15s), then 1ml of 1:1 hexane: ethyl acetate was added. Samples were vortexed for 5 mins then centrifuged at 16.1rfc for 5 mins. 1ml of supernatant was removed and evaporated to dryness under a stream of nitrogen. Samples were reconstituted with 50µl of mobile phase A. 10µl injections were made on the platform.

Analysis of the data was done using QuanLynxs software (Waters Corporation, Milford, MA, USA) by generating a linear equation based on peak area ratios (PAR) of the analyte over the internal standard peak area. This was then compiled against the expected concentration of the calibrators to generate a slope and intercept to be applied back to the PAR’s to obtain a return value. Acceptability of levels of quantitation was determined by a variance less than 15% from the expected return value.

For all studies a Water’s Xevo TQ-XS coupled to a Water’s I-Class Ultra high-pressure liquid chromatography system (Waters Corporation, Milford, MA, USA) was used for analysis: Instrument was operated in electrospray positive ionization mode (ESI-POS) with the following settings: Capillary voltage (kV) – 3.1, Desolvation Temp (C) – 350, Desolvation gas flow (L/Hr) – 800 nitrogen.

### Statistical analyses

All statistical analyses were performed using GraphPad Prism as detailed in figure legends. Unless otherwise stated, error bars represent standard error of them mean (SEM), and p<0.05 is considered statistically significant.

Unless otherwise stated, pairwise comparisons utilized the unpaired, unequal variances, t-test (Welch’s test).

### Scientific rigor

Sex – approximately equal numbers of male and female animals were used in each experiment. No sex differences have yet been reported in the *Ndufs4*(KO) mice.

Sample size determination and control distribution – all experiments were initially run with an n=4 animals per treatment group, which provides 80% power to detect changes of 30% at a p-value of 0.05, assuming standard deviation of 15% from the mean. Additional animals were added if variance was high, or trends toward significance were observed (to bolster or invalidate trending values). In general, untreated *Ndufs4*(KO) animals show high variance when disease is advanced, necessitating increased replicate numbers (such as in plethysmography analyses). All lifespan curves include at minimum 6 animals from different litters. Control treated *Ndufs4*(KO) groups are more heavily populated for three reasons: first, variance is highest in this group; second, increasing n in this group adds to statistical power of all treatment comparisons; third, control treated animals were included throughout the course of these studies to ensure not drift in the behavior of our colony as treatment groups were studied.

Randomization and blinding – all treatment groups were randomly assigned. Blinding was performed during image analysis. Full blinding of treatments was not possible due to hair color and body size changes induced by treatments, as detailed.

Inclusion/exclusion criteria – animals showing severe weaning stress, born as runts, or born with unrelated health issues (for example hydrocephaly) were excluded. Otherwise, no animals were excluded from randomization to treatment groups.

## Acknowledgments

Dedication: We dedicate this study to Alexandra Serex, Steve and Teresa Serex, their family, and all those living with the consequences of mitochondrial disease. They continue to inspire us.

National Institutes of Health grant NIH/GM R00 126147 (SCJ)

National Institutes of Health grant NIH/GM R01 133865 (MS, SCJ)

Northwest Mitochondrial Research Guild (SCJ)

North American Mitochondrial Disease Consortium (JS, SCJ)

National Institutes of Health T32 GM086270 (KS)

## Author Contributions

Conceptualization: Simon C Johnson

Methodology: Simon C Johnson, Julia Claire Stokes, Rebecca L Bornstein, Katerina James, Nathan Baertsch, Kyung Yeon Park, Kira Spencer

Investigation: Simon C Johnson, Julia Claire Stokes, Rebecca L Bornstein, Katerina James, Nathan Baertsch, Kyung Yeon Park, Kira Spencer, Brittany M Johnson, Katie Vo

Visualization: Simon C Johnson, Rebecca L Bornstein, Nathan Baertsch

Funding acquisition: Simon C Johnson, Margaret Sedensky

Project administration: Simon C Johnson

Supervision: Simon C Johnson, Margaret M Sedensky, Philip G Morgan

Writing – original draft: Simon C Johnson

Writing – review & editing: Simon C Johnson, Margaret M Sedensky, Philip G Morgan, Rebecca L Bornstein

## Data and materials availability

All data are available in the main text or the supplementary materials.

## Figures

**Fig. S1.**
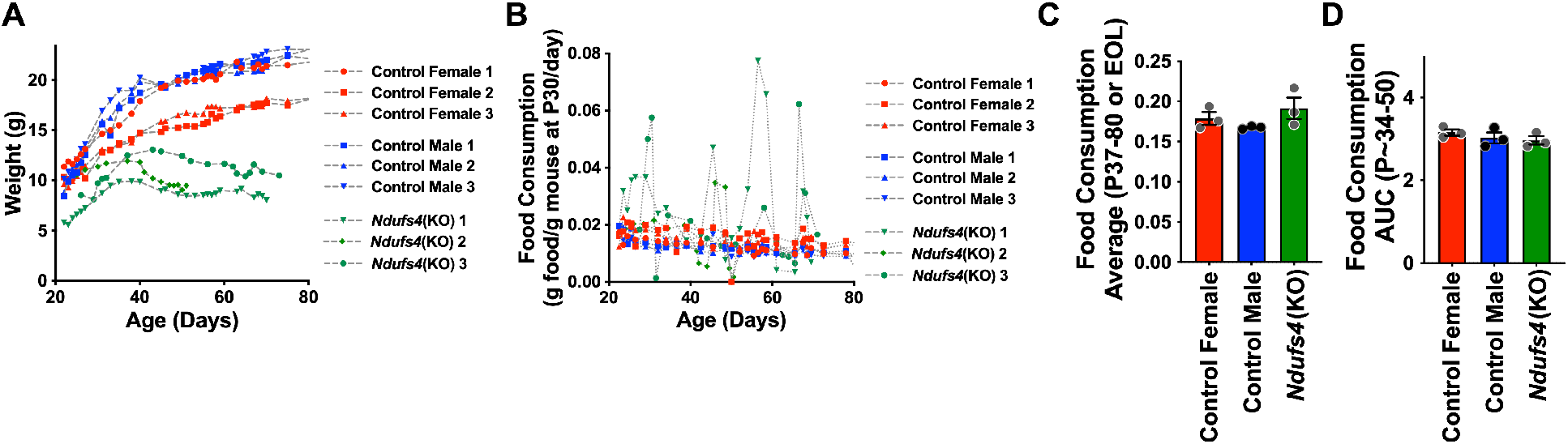
Cachexia occurs without anorexia in *Ndufs4*(KO). (A) Daily weight and (B) food consumption measurements for control and *Ndufs4*(KO) animals. *Ndufs4*(KO) mice are small compared to control animals, so consumption is normalized to weight at P30, a post-weaning, but pre-disease onset, weight. While there was a noticeable presence of day-to-day food consumption outlier days in the *Ndufs4*(KO) mice (B), neither the average food consumption per day (A) or total food consumption (B) throughout the period from onset of cachexia to death were different in the *Ndufs4*(KO) animals when normalizing to animal weight at P30. The age-range in (D) was set to span the period shortly prior to the onset of cachexia and continue until the first mouse was lost to euthanasia.

**Fig. S2.**
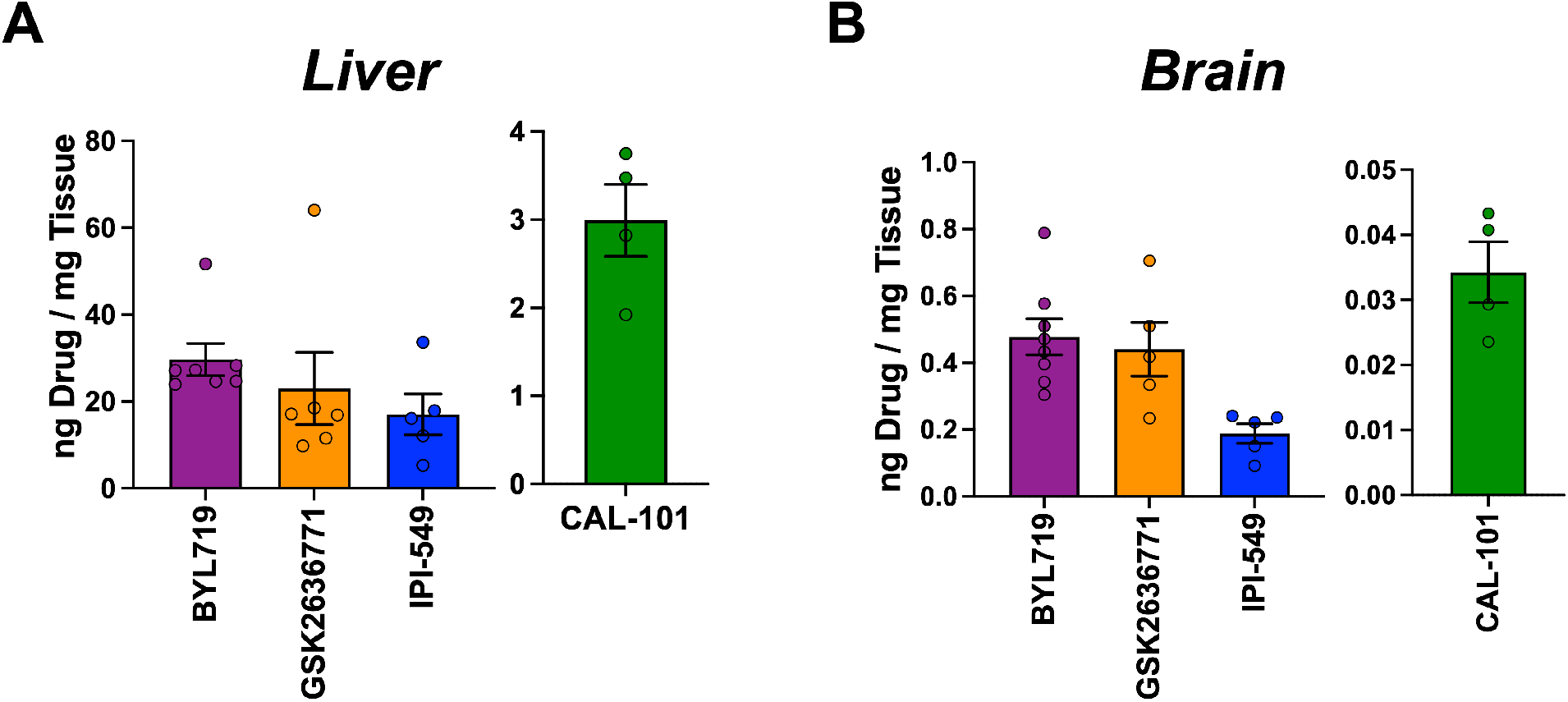
Tissue levels of PI3K p110 catalytic subunit isoform specific inhibitors. (A) Liver and (B) brain tissue levels of BYL719, GSK2636771, IPI-549, and CAL-101 (see ***Methods***).

**Fig S3.**
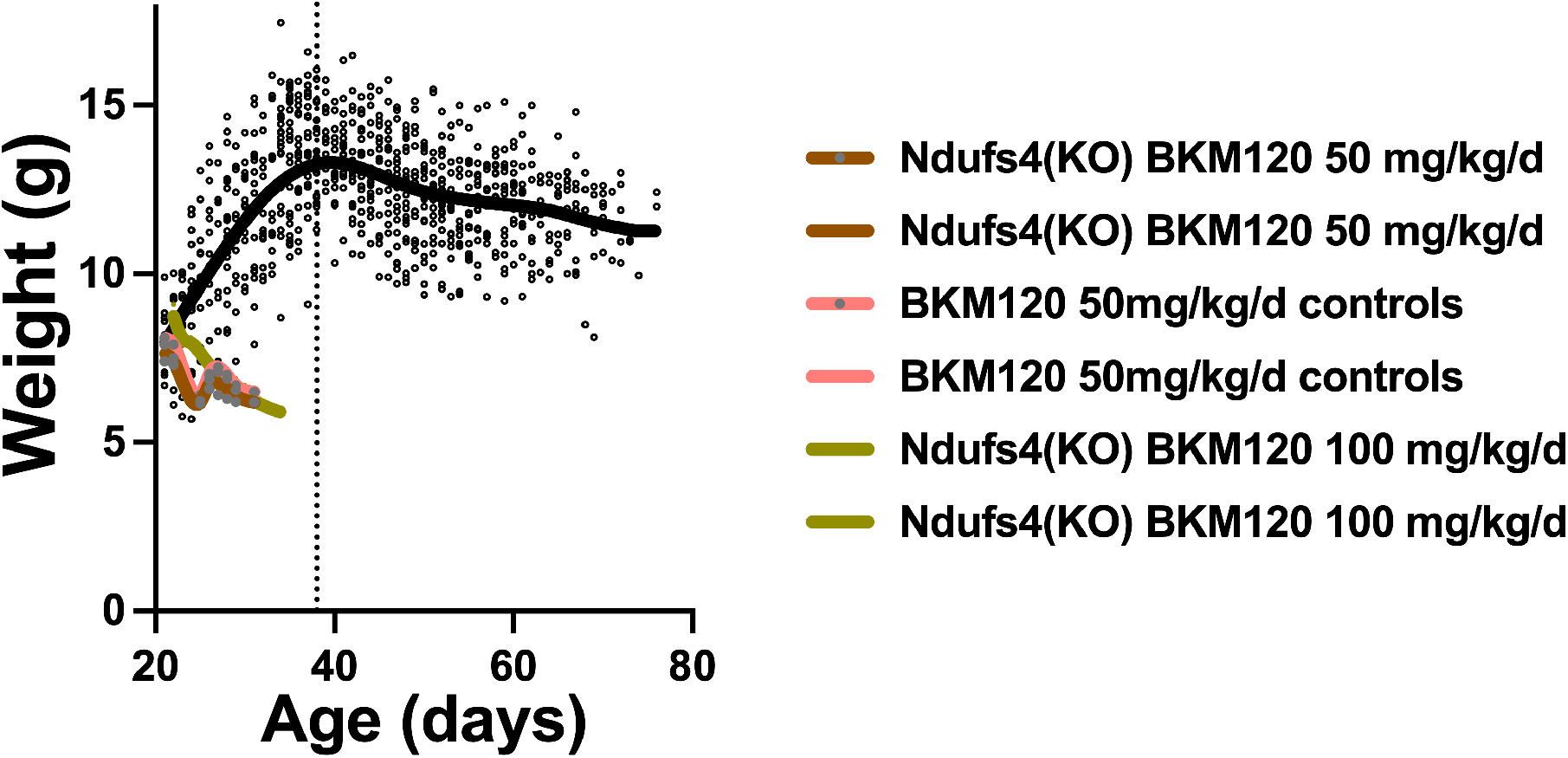
Impact of BKM120 during post-weaning development. BKM120 treatment at 50 or 100 mg/kg/day led to weight loss and euthanasia in both control and *Ndufs4*(KO) mice when treated during post-weaning development. N=2 animals per dose and genotype for BKM120.

**Fig. S4.**
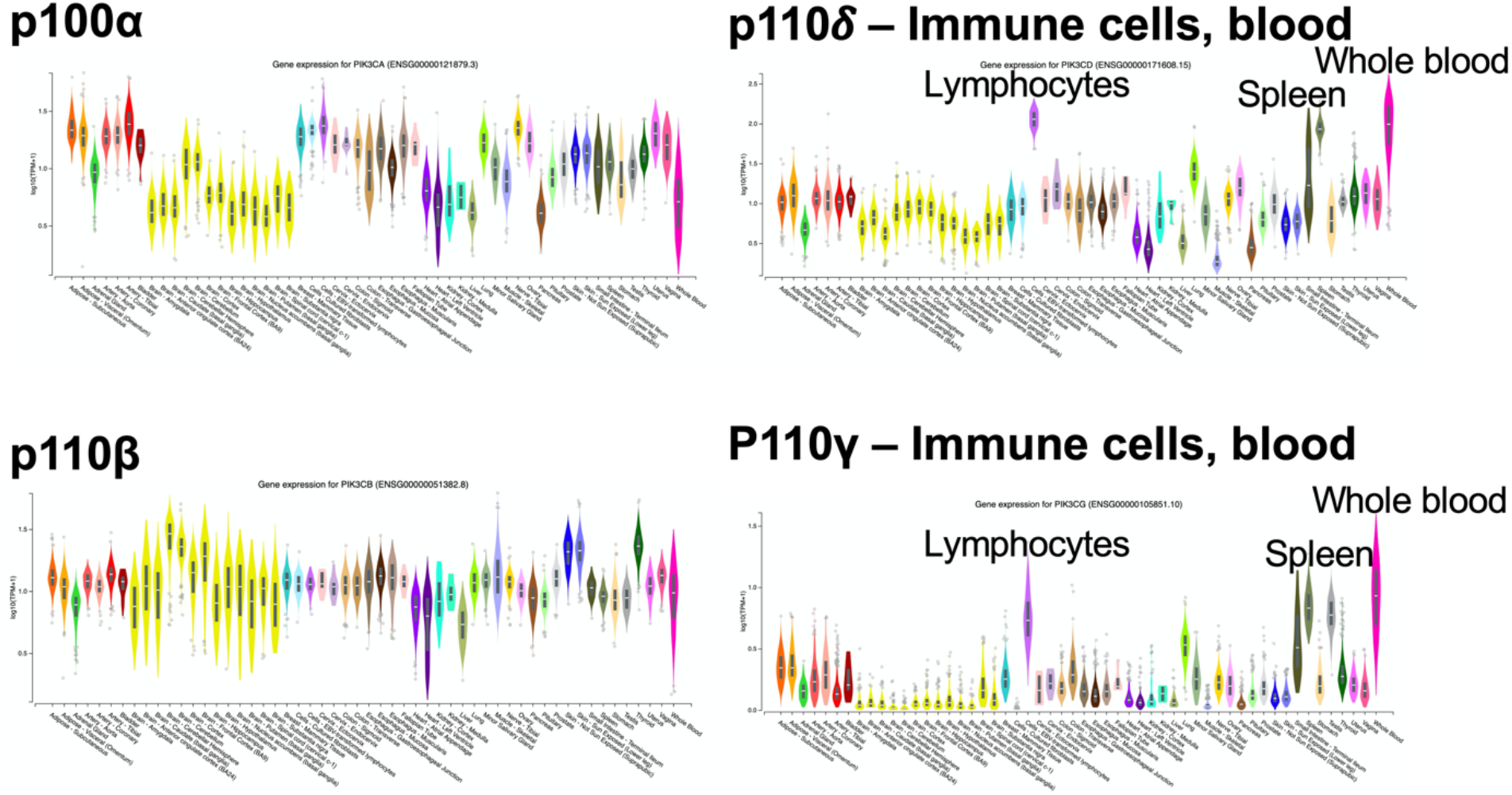
Expression pattern of PI3K p110 catalytic subunit isoforms. Expression of p110α, p110β, p110δ, and p110γ isoforms in different human tissues. Data and figures collected from the GTEx database by searching for PIK3CA, PIK3CB, PIK3CG, and PIK3CD. https://gtexportal.org/home/gene/

**Fig. S5.**
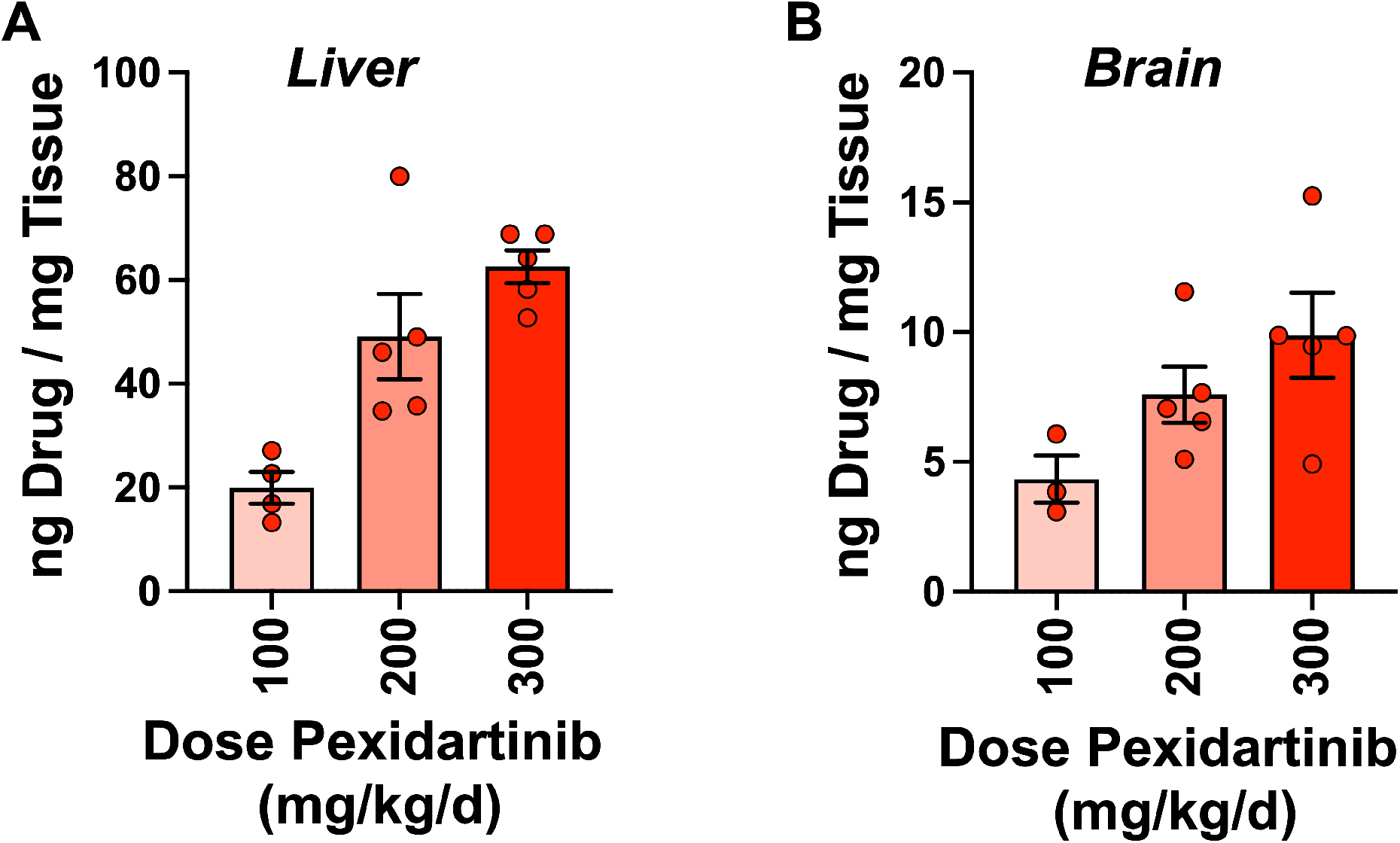
Tissue levels of drug in mice treated with 100, 200, or 300 mg/kg/day pexidartinib in chow. (A) Liver and (B) brain drug levels. See ***Methods***.

**Fig. S6.**
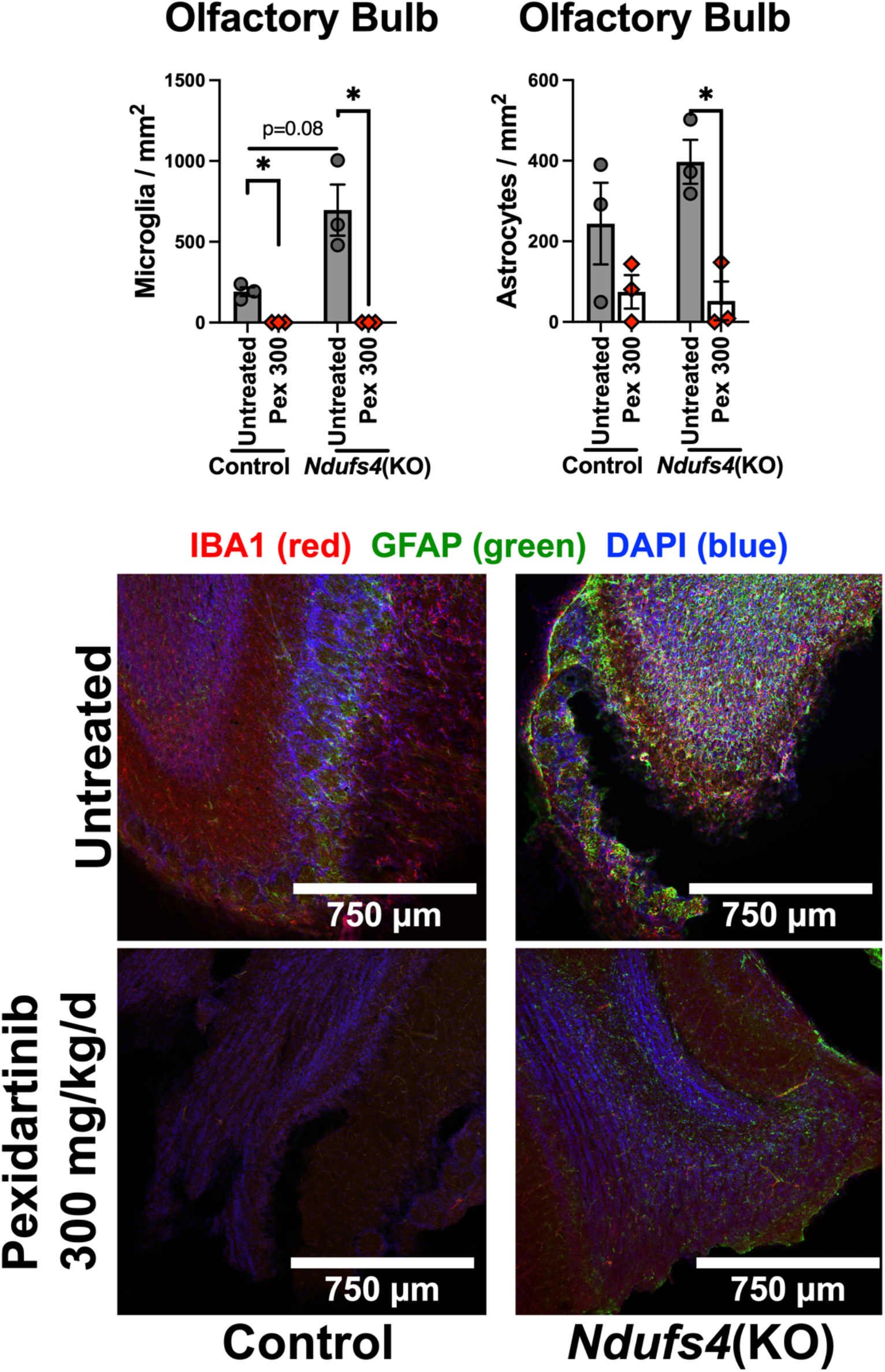
Impact of pexidartinib on olfactory bulb microgliosis and astrocytosis. Pexidartinib treatment appears to rescue neuroinflammation in the olfactory bulb, though high variance and limited sample numbers preclude statistical significance in some comparisons. Representative images shown.

**Fig. S7.**
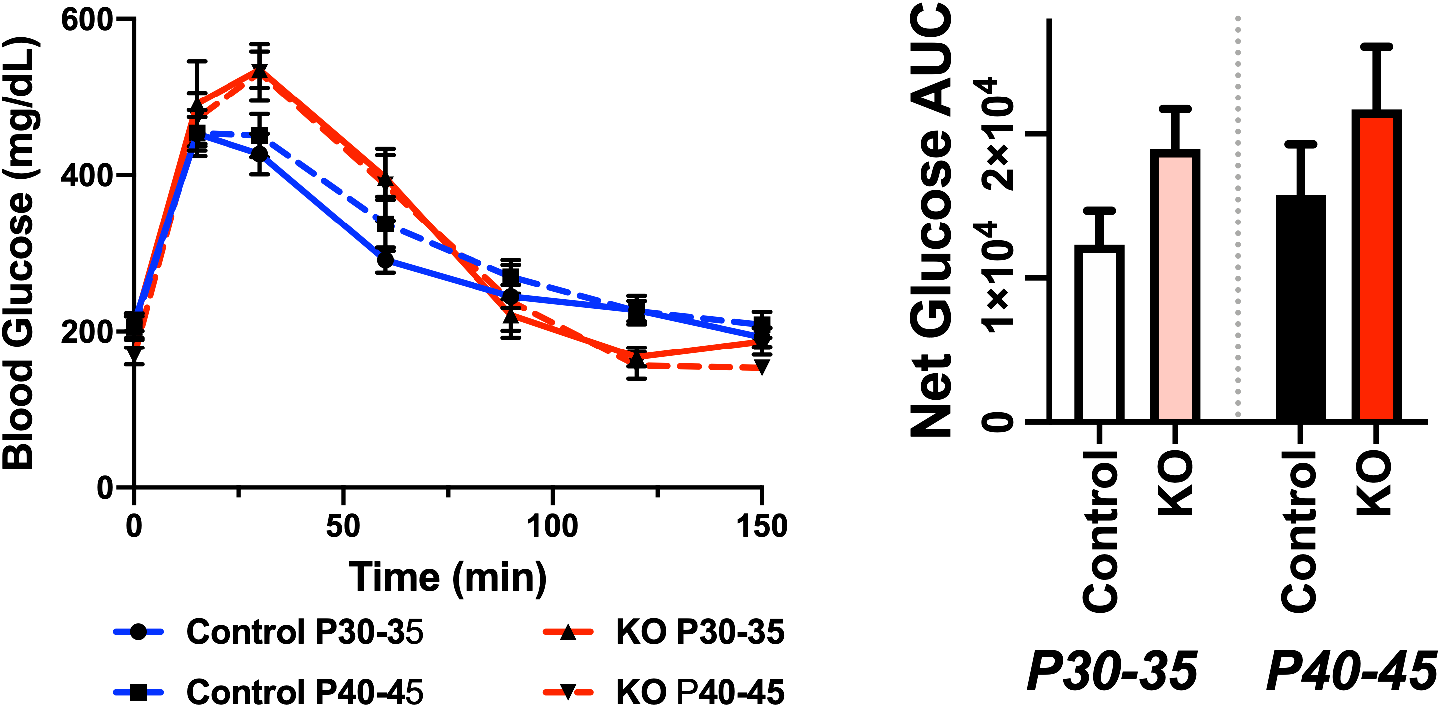
Glucose clearance in glucose tolerance test. Glucose clearance in the GTT assay does not change as a function of age from pre-to post-P37. Area under the curve (AUC) for glucose trends up in *Ndufs4*(KO) animals at both ages but is non-significant and the difference does not change with age. N=10 control animals per age, 4 *Ndufs4*(KO) at P30-35 and 8 at P40-45.

## References

1. F. G. Goncalves et al., Primary Mitochondrial Disorders of the Pediatric Central Nervous System: Neuroimaging Findings. Radiographics 40, 2042–2067 (2020)

2. C. L. Alston, S. L. Stenton, G. Hudson, H. Prokisch, R. W. Taylor, The genetics of mitochondrial disease: dissecting mitochondrial pathology using multi-omic pipelines. J Pathol 254, 430–442 (2021)

3. N. J. Lake, M. J. Bird, P. Isohanni, A. Paetau, Leigh syndrome: neuropathology and pathogenesis. J Neuropathol Exp Neurol 74, 482–492 (2015)

4. S. C. Johnson et al., Dose-dependent effects of mTOR inhibition on weight and mitochondrial disease in mice. Front Genet 6, 247 (2015).PMC4510413

5. S. C. Johnson et al., mTOR inhibition alleviates mitochondrial disease in a mouse model of Leigh syndrome. Science 342, 1524–1528 (2013).PMC4055856

6. S. C. Johnson et al., mTOR inhibitors may benefit kidney transplant recipients with mitochondrial diseases. Kidney Int 95, 455–466 (2019).PMC6389356

7. A. Sage-Schwaede et al., Exploring mTOR inhibition as treatment for mitochondrial disease. Ann Clin Transl Neurol 6, 1877–1881 (2019).PMC6764630

8. A. Saudemont et al., p110gamma and p110delta isoforms of phosphoinositide 3-kinase differentially regulate natural killer cell migration in health and disease. Proc Natl Acad Sci U S A 106, 5795–5800 (2009).PMC2667007

9. C. Fritsch et al., Characterization of the novel and specific PI3Kalpha inhibitor NVP-BYL719 and development of the patient stratification strategy for clinical trials. Mol Cancer Ther 13, 1117–1129 (2014)

10. C. Leroy et al., Activation of IGF1R/p110beta/AKT/mTOR confers resistance to alpha-specific PI3K inhibition. Breast Cancer Res 18, 41 (2016).PMC4820873

11. K. J. Pridham et al., PIK3CB/p110beta is a selective survival factor for glioblastoma. Neuro Oncol 20, 494–505 (2018).PMC5909664

12. R. J. Davis et al., Anti-PD-L1 Efficacy Can Be Enhanced by Inhibition of Myeloid-Derived Suppressor Cells with a Selective Inhibitor of PI3Kdelta/gamma. Cancer Res 77, 2607–2619 (2017).PMC5466078

13. M. T. Burger et al., Identification of NVP-BKM120 as a Potent, Selective, Orally Bioavailable Class I PI3 Kinase Inhibitor for Treating Cancer. ACS Med Chem Lett 2, 774–779 (2011).PMC4017971

14. M. Schubert Baldo, L. Vilarinho, Molecular basis of Leigh syndrome: a current look. Orphanet J Rare Dis 15, 31 (2020).PMC6990539

15. A. M. Gonzalez-Angulo et al., Weekly nab-Rapamycin in patients with advanced nonhematologic malignancies: final results of a phase I trial. Clin Cancer Res 19, 5474–5484 (2013).PMC3935482

16. W. A. Denny, J. U. Flanagan, Small-molecule CSF1R kinase inhibitors; review of patents 2015-present. Expert Opin Ther Pat 31, 107–117 (2021)

17. A. Quintana et al., Fatal breathing dysfunction in a mouse model of Leigh syndrome. J Clin Invest 122, 2359–2368 (2012).PMC3387817

18. S. C. Ramirez et al., Perinatal Breathing Patterns and Survival in Mice Born Prematurely and at Term. Front Physiol 10, 1113 (2019).PMC6728753

19. H. F. Lee, C. S. Chi, C. R. Tsai, C. H. Chen, Epileptic seizures in infants and children with mitochondrial diseases. Pediatr Neurol 45, 169–174 (2011)

20. J. M. Cameron et al., Exome sequencing identifies complex I NDUFV2 mutations as a novel cause of Leigh syndrome. Eur J Paediatr Neurol 19, 525–532 (2015)

21. S. Fukumura et al., Compound heterozygous GFM2 mutations with Leigh syndrome complicated by arthrogryposis multiplex congenita. J Hum Genet 60, 509–513 (2015)

22. J. Yokoyama et al., [A case of rhabdomyolysis after status epilepticus without stroke-like episodes in mitochondrial myopathy, encephalopathy, lactic acidosis, and stroke-like episodes]. Rinsho Shinkeigaku 56, 204–207 (2016)

23. K. Yamada et al., Diagnostic accuracy of blood and CSF lactate in identifying children with mitochondrial diseases affecting the central nervous system. Brain Dev 34, 92–97 (2012)

24. C. S. Chi et al., Lactate peak on brain MRS in children with syndromic mitochondrial diseases. J Chin Med Assoc 74, 305–309 (2011)

25. Y. Takahashi et al., Detection of increased intracerebral lactate in a mouse model of Leigh syndrome using proton MR spectroscopy. Magn Reson Imaging 58, 38–43 (2019)

26. M. Nafisinia et al., Whole Exome Sequencing Identifies the Genetic Basis of Late-Onset Leigh Syndrome in a Patient with MRI but Little Biochemical Evidence of a Mitochondrial Disorder. JIMD Rep 32, 117–124 (2017).PMC5362551

27. P. A. Kramer, S. Ravi, B. Chacko, M. S. Johnson, V. M. Darley-Usmar, A review of the mitochondrial and glycolytic metabolism in human platelets and leukocytes: implications for their use as bioenergetic biomarkers. Redox Biol 2, 206–210 (2014).PMC3909784

28. V. C. Hsieh, J. Niezgoda, M. M. Sedensky, C. L. Hoppel, P. G. Morgan, Anesthetic Hypersensitivity in a Case-Controlled Series of Patients With Mitochondrial Disease. Anesth Analg, (2021).PMC8280249

29. E. B. Kayser, P. G. Morgan, M. M. Sedensky, GAS-1: a mitochondrial protein controls sensitivity to volatile anesthetics in the nematode Caenorhabditis elegans. Anesthesiology 90, 545–554 (1999)

30. J. M. Sonner, D. Gong, E. I. Eger, 2nd, Naturally occurring variability in anesthetic potency among inbred mouse strains. Anesth Analg 91, 720–726 (2000)

31. J. H. Lewis et al., Pexidartinib Long-Term Hepatic Safety Profile in Patients with Tenosynovial Giant Cell Tumors. Oncologist 26, e863–e873 (2021).PMC8100574

32. H. Ikeda et al., PI3K/p110{delta} is a novel therapeutic target in multiple myeloma. Blood 116, 1460–1468 (2010).PMC2938837

33. L. S. Carnevalli et al., PI3Kalpha/delta inhibition promotes anti-tumor immunity through direct enhancement of effector CD8(+) T-cell activity. J Immunother Cancer 6, 158 (2018).PMC6307194

34. D. Leigh, Subacute necrotizing encephalomyelopathy in an infant. J Neurol Neurosurg Psychiatry 14, 216–221 (1951).PMC499520

35. A. Quintana, S. E. Kruse, R. P. Kapur, E. Sanz, R. D. Palmiter, Complex I deficiency due to loss of Ndufs4 in the brain results in progressive encephalopathy resembling Leigh syndrome. Proc Natl Acad Sci U S A 107, 10996–11001 (2010).PMC2890717

36. C. Kohler, Allograft inflammatory factor-1/Ionized calcium-binding adapter molecule 1 is specifically expressed by most subpopulations of macrophages and spermatids in testis. Cell Tissue Res 330, 291–302 (2007)

37. F. Pierezan, J. Mansell, A. Ambrus, A. Rodrigues Hoffmann, Immunohistochemical expression of ionized calcium binding adapter molecule 1 in cutaneous histiocytic proliferative, neoplastic and inflammatory disorders of dogs and cats. J Comp Pathol 151, 347–351 (2014)

38. T. A. van Wageningen et al., Regulation of microglial TMEM119 and P2RY12 immunoreactivity in multiple sclerosis white and grey matter lesions is dependent on their inflammatory environment. Acta Neuropathol Commun 7, 206 (2019).PMC6907356

39. S. C. Johnson, Translational Medicine. A target for pharmacological intervention in an untreatable human disease. Science 346, 1192 (2014)

40. S. E. Siegmund et al., Low-dose rapamycin extends lifespan in a mouse model of mtDNA depletion syndrome. Hum Mol Genet 26, 4588–4605 (2017).PMC5886265

41. G. Civiletto et al., Rapamycin rescues mitochondrial myopathy via coordinated activation of autophagy and lysosomal biogenesis. EMBO Mol Med 10, (2018).PMC6220341

42. L. Wischhof et al., A disease-associated Aifm1 variant induces severe myopathy in knockin mice. Mol Metab 13, 10–23 (2018).PMC6026322

43. A. Wang, J. Mouser, J. Pitt, D. Promislow, M. Kaeberlein, Rapamycin enhances survival in a Drosophila model of mitochondrial disease. Oncotarget 7, 80131–80139 (2016).PMC5348310

44. F. Fan, R. Sam, E. Ryan, K. Alvarado, E. Villa-Cuesta, Rapamycin as a potential treatment for succinate dehydrogenase deficiency. Heliyon 5, e01217 (2019).PMC6374580

45. S. C. Johnson et al., Regional metabolic signatures in the Ndufs4(KO) mouse brain implicate defective glutamate/alpha-ketoglutarate metabolism in mitochondrial disease. Mol Genet Metab 130, 118–132 (2020).PMC7272141

46. T. K. Ito et al., Hepatic S6K1 Partially Regulates Lifespan of Mice with Mitochondrial Complex I Deficiency. Front Genet 8, 113 (2017).PMC5585733

47. C. F. Lee, A. Caudal, L. Abell, G. A. Nagana Gowda, R. Tian, Targeting NAD(+) Metabolism as Interventions for Mitochondrial Disease. Sci Rep 9, 3073 (2019).PMC6395802

48. M. Martin-Perez et al., PKC downregulation upon rapamycin treatment attenuates mitochondrial disease. Nat Metab 2, 1472–1481 (2020).PMC8017771

49. G. S. McElroy et al., NAD+ Regeneration Rescues Lifespan, but Not Ataxia, in a Mouse Model of Brain Mitochondrial Complex I Dysfunction. Cell Metab 32, 301–308 e306 (2020).PMC7415718

50. I. H. Jain et al., Leigh Syndrome Mouse Model Can Be Rescued by Interventions that Normalize Brain Hyperoxia, but Not HIF Activation. Cell Metab 30, 824–832 e823 (2019)

51. I. H. Jain et al., Hypoxia as a therapy for mitochondrial disease. Science 352, 54–61 (2016).PMC4860742

52. I. Bolea et al., Defined neuronal populations drive fatal phenotype in a mouse model of Leigh syndrome. Elife 8, (2019).PMC6731060

53. G. Karamanlidis et al., Mitochondrial complex I deficiency increases protein acetylation and accelerates heart failure. Cell Metab 18, 239–250 (2013).PMC3779647

54. H. Zhang et al., Heart specific knockout of Ndufs4 ameliorates ischemia reperfusion injury. J Mol Cell Cardiol 123, 38–45 (2018).PMC6192835

55. Y. A. Chiao et al., NAD(+) Redox Imbalance in the Heart Exacerbates Diabetic Cardiomyopathy. Circ Heart Fail 14, e008170 (2021).PMC8373812

56. J. L. Edmonds et al., The otolaryngological manifestations of mitochondrial disease and the risk of neurodegeneration with infection. Arch Otolaryngol Head Neck Surg 128, 355–362 (2002)

57. Y. J. Lee, S. K. Hwang, S. Kwon, Acute Necrotizing Encephalopathy in Children: a Long Way to Go. J Korean Med Sci 34, e143 (2019).PMC6522889

58. D. M. Niyazov, S. G. Kahler, R. E. Frye, Primary Mitochondrial Disease and Secondary Mitochondrial Dysfunction: Importance of Distinction for Diagnosis and Treatment. Mol Syndromol 7, 122–137 (2016).PMC4988248

59. F. Porta et al., SLC25A19 deficiency and bilateral striatal necrosis with polyneuropathy: a new case and review of the literature. J Pediatr Endocrinol Metab 34, 261–266 (2021)

60. H. S. Wang, S. C. Huang, Acute necrotizing encephalopathy of childhood. Chang Gung Med J 24, 1–10 (2001)

61. Y. Wei, L. Wang, Adult-onset Leigh syndrome with central fever and peripheral neuropathy due to mitochondrial 9176T>C mutation. Neurol Sci 39, 2225–2228 (2018)

62. T. S. Wu et al., Leigh disease (subacute necrotizing encephalomyelopathy): report of one case. Zhonghua Min Guo Xiao Er Ke Yi Xue Hui Za Zhi 34, 301–307 (1993)

63. H. J. S. Dawkins et al., Progress in Rare Diseases Research 2010-2016: An IRDiRC Perspective. Clin Transl Sci 11, 11–20 (2018).PMC5759730

64. R. Gopal-Srivastava, P. Kaufmann, Facilitating Clinical Studies in Rare Diseases. Adv Exp Med Biol 1031, 125–140 (2017)

65. P. Kaufmann, A. R. Pariser, C. Austin, From scientific discovery to treatments for rare diseases -the view from the National Center for Advancing Translational Sciences - Office of Rare Diseases Research. Orphanet J Rare Dis 13, 196 (2018).PMC6219030

66. A. Rath et al., A systematic literature review of evidence-based clinical practice for rare diseases: what are the perceived and real barriers for improving the evidence and how can they be overcomeã Trials 18, 556 (2017).PMC5700662

67. X. Chang et al., A meta-analysis and systematic review of Leigh syndrome: clinical manifestations, respiratory chain enzyme complex deficiency, and gene mutations. Medicine (Baltimore) 99, e18634 (2020).PMC7004636

68. R. D. S. Pitceathly, N. Keshavan, J. Rahman, S. Rahman, Moving towards clinical trials for mitochondrial diseases. J Inherit Metab Dis 44, 22–41 (2021).PMC8432143

69. J. S. Lee et al., Genetic heterogeneity in Leigh syndrome: Highlighting treatable and novel genetic causes. Clin Genet 97, 586–594 (2020)

70. S. M. Davis et al., Leukemia inhibitory factor modulates the peripheral immune response in a rat model of emergent large vessel occlusion. J Neuroinflammation 15, 288 (2018).PMC6190542

71. X. Zeng et al., Interferon-inducible protein 10, but not monokine induced by gamma interferon, promotes protective type 1 immunity in murine Klebsiella pneumoniae pneumonia. Infect Immun 73, 8226–8236 (2005).PMC1307052

72. V. K. Kommineni, C. N. Nagineni, A. William, B. Detrick, J. J. Hooks, IFN-gamma acts as anti-angiogenic cytokine in the human cornea by regulating the expression of VEGF-A and sVEGF-R1. Biochem Biophys Res Commun 374, 479–484 (2008).PMC2997485

73. D. Bastian, Y. Wu, B. C. Betts, X. Z. Yu, The IL-12 Cytokine and Receptor Family in Graft-vs.-Host Disease. Front Immunol 10, 988 (2019).PMC6518430

74. M. Yabe et al., Role of interleukin-12 in the development of acute graft-versus-host disease in bone marrow transplant patients. Bone Marrow Transplant 24, 29–34 (1999)

75. B. R. Blazar, P. A. Taylor, A. Panoskaltsis-Mortari, D. A. Vallera, Rapamycin inhibits the generation of graft-versus-host disease- and graft-versus-leukemia-causing T cells by interfering with the production of Th1 or Th1 cytotoxic cytokines. J Immunol 160, 5355–5365 (1998)

76. I. Couillin, N. Riteau, STING Signaling and Sterile Inflammation. Front Immunol 12, 753789 (2021).PMC8517477

77. Y. Zhu et al., STING: a master regulator in the cancer-immunity cycle. Mol Cancer 18, 152 (2019).PMC6827255

78. J. Stokes et al., Mechanisms underlying neonate-specific metabolic effects of volatile anesthetics. Elife 10, (2021).PMC8291971

